# From farm to fork: persistence of clinically-relevant multidrug-resistant and copper-tolerant *Klebsiella pneumoniae* long after colistin withdrawal in poultry production

**DOI:** 10.1101/2023.04.03.535403

**Authors:** Joana Mourão, Marisa Ribeiro-Almeida, Carla Novais, Mafalda Magalhães, Andreia Rebelo, Sofia Ribeiro, Luísa Peixe, Ângela Novais, Patrícia Antunes

**Affiliations:** UCIBIO – Applied Molecular Biosciences Unit, REQUIMTE, Laboratory of Microbiology, Department of Biological Sciences, Faculty of Pharmacy, University of Porto, Portugal; Associate Laboratory i4HB - Institute for Health and Bioeconomy, Faculty of Pharmacy, University of Porto, Porto, Portugal; Center for Innovative Biomedicine and Biotechnology (CIBB), University of Coimbra, Coimbra, Portugal; School of Medicine and Biomedical Sciences, University of Porto (ICBAS-UP), Porto, Portugal; Faculty of Nutrition and Food Sciences, University of Porto, Porto, Portugal; ESS, Polytechnic of Porto, Porto, Portugal

**Author notes:** Address correspondence to Patrícia Antunes. Joana Mourão, Carla Novais, Luísa Peixe and Patrícia Antunes are active members of the European Society of Clinical Microbiology and Infectious Diseases (ESCMID) Food and Water-borne Infection Study Group (EFWISG).

**Keywords:** chicken meat, poultry sector, copper-feed supplementation, antibiotic resistance, IncF plasmids, One Health, food safety.

## Abstract

The concern of colistin-resistant bacteria in animal-food-environmental-human ecosystems prompted the poultry sector to implement colistin restrictions and explore alternative trace metals/copper feed supplementation. The impact of these strategies on the selection and persistence of colistin-resistant *Klebsiella pneumoniae* (Kp) in the whole poultry-production chain needs clarification. We assessed colistin-resistant and copper-tolerant Kp occurrence in chicken raised with inorganic and organic copper-formulas from one-day-old chicks to meat (7 farms/2019-2020), after long-term colistin withdrawal (>2-years). Clonal diversity and Kp adaptive features were characterized by cultural, molecular, and whole-genome-sequencing (WGS) approaches. Most chicken-flocks (75%) carried Kp at early+pre-slaughter stages, with a significant decrease (p<0.05) in meat batches (17%) and sporadic water/feed contamination. High rates (>50%) of colistin-resistant/*mcr*-negative Kp were observed among faecal samples, independently of feed. Most samples carried multidrug-resistant (90%) and copper-tolerant isolates (81%; *pco+sil*/MIC_CuSO4_ ≥16mM). WGS revealed accumulation of colistin resistance associated mutations and F-type multireplicon plasmids carrying antibiotic resistance and metal/copper-tolerance genes. The Kp population was polyclonal, with various lineages dispersed throughout poultry production. ST15-KL19, ST15-KL146 and ST392-KL27, and IncF plasmids were similar to those from global human clinical isolates, suggesting chicken-production as a reservoir/source of clinically-relevant Kp lineages and genes with potential risk to humans through food and/or environmental exposure. Despite long-term colistin ban limited *mcr* spread, it was ineffective in controlling colistin-resistant/*mcr*-negative Kp, regardless of feed. This study provides crucial insights into the persistence of clinically-relevant Kp in the poultry-production chain and highlights the need for continued surveillance and proactive food safety actions within a ’One-Health’ perspective.

**IMPORTANCE:** The spread of bacteria resistant to last-resort antibiotics such as colistin throughout the food chain is a serious concern for public health. The poultry sector has responded by restricting colistin use and exploring alternative trace metals/copper feed supplements. However, it is unclear how and to which extent these changes impact the selection and persistence of clinically-relevant *Klebsiella pneumoniae* (Kp) throughout poultry chain. We found a high occurrence of copper-tolerant and colistin-resistant/*mcr*-negative Kp in chicken flocks, regardless of inorganic and organic copper-formulas and long-term colistin ban. Despite the high Kp diversity, the occurrence of identical lineages and plasmids across samples and/or clinical isolates suggests poultry as a potential source of human Kp exposure. This study highlights the need for continued surveillance and proactive farm-to-fork actions to mitigate the risks to public health, relevant for stakeholders involved in food industry and policymakers tasked with regulating food safety.

## INTRODUCTION

A major public health concern has been attributed to colistin-resistant bacteria, particularly those carrying mobilized colistin resistance (*mcr*) genes, due to implications for human, animal and environmental health (1, 2). The wide use of colistin for decades in veterinary medicine imposed global restrictions on the food-animal production (3–5). In the EU, colistin is currently classified in the *category B - Restrict* (only for clinical infections in the absence of an alternative) in line with applicable Regulations from 2022 (prohibiting all routine antibiotic use in farming) and the 2030 European goal (to reduce 50% the sales of antibiotics for farm animals), with impact in different environmental compartments (6–10). Diverse studies revealed a decreasing occurrence of *mcr*-carrying bacteria (mainly *Escherichia coli* and *Klebsiella pneumoniae*) in farms shortly after colistin restrictions (11, 12), whereas the persistence of colistin-resistant bacteria by mechanisms other than *mcr* is unknown. Furthermore, there are no studies evaluating the long-term impact of colistin restriction on the occurrence and diversity of colistin-resistant bacteria throughout food-animal production environments and/or considering farm-level variation or food-chain practices (4, 11). Studies on intensive poultry meat systems are ideally suited to inform reducing/replacing strategies since the EU is one of the top four global chicken-meat producers with an upward trend in consumer preferences for "greener" antibiotic-free food products (13–15).

In the same line as colistin restriction, is the development of nutritional alternatives to improve poultry health and growth, including the use of copper (Cu) in the feeds (3, 15–17). Copper is a heavy metal widely used in poultry feed due to its requirement in many physiologic processes, immune-stimulating actions, and antimicrobial activity in the gut (15, 18, 19). In current commercial practice, inorganic trace minerals formulation feeds (ITMF) are supplemented with inorganic sources of Cu (e.g., Cu sulphate, Cu oxide) in high levels (<25 mg/kg, accordingly to EU Regulation 2018/1039), leading to potential environmental accumulation (20–22). Alternatively, organic trace minerals feed supplements (OTMF) (e.g., Cu chelated with amino acids, peptides, or proteins) have been associated with a higher Cu bioavailability in the digestive tract than ITMF and an animal performance improvement, being expected a reduced faecal excretion and environmental release (19–23). However, the effect of different Cu-supplemented feeds on gastrointestinal microbiota composition is still unclear and needs to be further explored by experimental studies (18–20, 24). In addition, a link has been proposed between feed supplementation with Cu (and other metals) and the selection of antibiotic-resistant bacteria (24, 25). However, the impact of variable feed formulations (OTMF *versus* ITMF) in poultry gut bacteria, particularly in species often associated with colistin resistance, and throughout the whole food production chain, has been poorly explored.

*K. pneumoniae* has been frequently associated with colistin resistance and *mcr* carriage in the clinical setting but its occurrence in the animal-food-environmental interface is underexplored (26). Besides we have previously shown the co-localization of metal/Cu operons with the *mcr-1* gene in plasmids from *K. pneumoniae* isolated from chicken meat batches shortly after the colistin withdrawal (12), suggesting a Cu co-selection effect for the persistence of colistin resistance. In fact, *K. pneumoniae* is a versatile species with a natural ability to acquire antibiotic and metal resistance genes, enabling their survival and circulation among animal-environment-human ecosystems (27–29). However, it remains unclear how and to which extent restriction/replacement of colistin interventions in poultry production impacts the selection and persistence of colistin-resistant *K. pneumoniae* (ColR-Kp) and, eventually, clones with a potential risk of foodborne and/or environmental transmission.

We used a One-Health approach by spanning the poultry production chain (from one-day-old chicks to meat) to better understand the impact of long-term colistin ban on the occurrence of colistin-resistant and Cu-tolerant *K. pneumoniae,* and its association with ITMF and OTMF feed formulations. Besides, we applied whole genome sequencing (WGS) to characterize these isolates in terms of clonal diversity, antibiotic resistance, metal tolerance and virulence determinants to have a comprehensive overview of the risks posed by the poultry production environment.

## RESULTS

### Occurrence of *K. pneumoniae* and colistin-resistant *K. pneumoniae* by samples

Our cultural approach detected *K. pneumoniae* in 42% (n=35/84) of the total tested samples within the poultry production chain. A high occurrence was found among faecal samples (82%; 78%-14/18 flocks at P1 and 88%-14/16 flocks at P2), corresponding to all but two of the flocks and all farms tested. However, a significant decrease in the occurrence was detected between P2 and chicken meat batches (17%-3/18 at P3, corresponding to flocks from 3 farms) (p<0,05) (**Fig. 1-A**). Only two flocks (2 farms) had *K. pneumoniae* isolates in all three stages (P1+P2+P3), while the remaining ones were found in P1+P2 (10 flocks) (75%-12/16 flocks at P1+P2±P3) or only in P1 (1 flock), P2 (2 flocks) or P3 (1 flock). Similar occurrence rates were observed between OTMF and ITMF samples in each of the three sampling stages (p>0,05) (**Fig. 1-A**). Regarding poultry house environmental samples, three feeds (2 farms) and one water sample (another farm) were contaminated with *K. pneumoniae*.

**Fig. 1.**
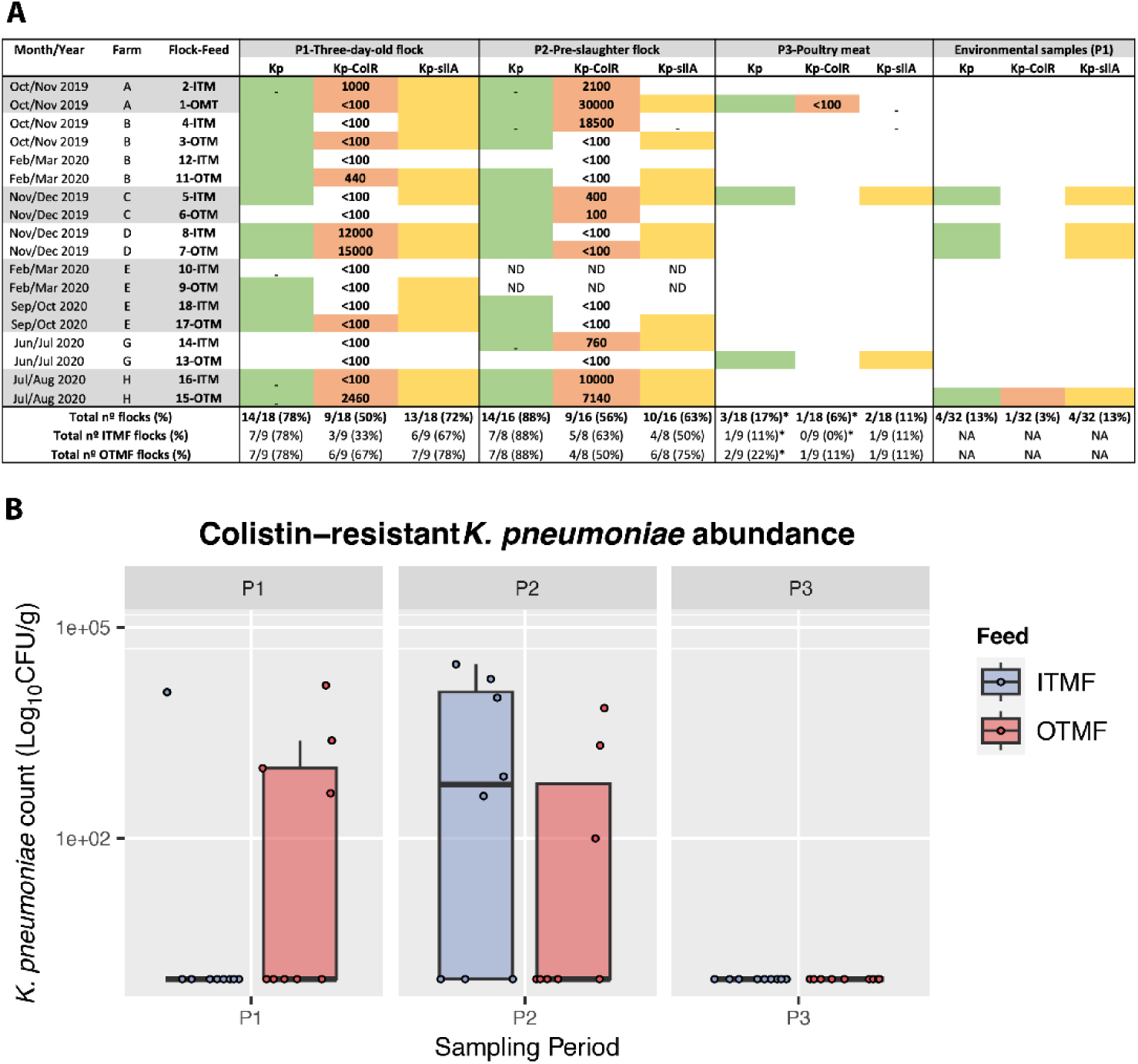
(**A)** Occurrence of *K. pneumoniae* (Kp), colistin-resistant *K. pneumoniae* (Kp-ColR) and *silA*-carrying *K. pneumoniae* (Kp-silA) among flocks (faeces, P1 and P2) and chicken meat (P3) by type of feed (ITMF or OTMF). Environmental samples 5, 7 and 8 were from feed and sample 15 corresponds to water. *, P<0,05 (Fisher exact test), when comparing P2 with P3. **(B)** Box plot of colistin-resistant *K. pneumoniae* counts among flocks (faeces, P1 and P2) and chicken meat (P3) by type of feed (ITMF or OTMF). P>0,05 was calculated using the Wilcoxon test for comparing P1 with P2 (both feeds). The median is represented by the black horizontal line and blue (ITMF) or red (OTMF) data points represent the counts in each sample.

The ColR-Kp isolates were recovered from all farms and sample types, corresponding to 57% (20/35) of *K. pneumoniae*-positive samples. Most were faecal samples at P1 (50%-9/18 flocks; 5 farms) and P2 (56%-9/16 flocks; 6 farms) stages, with no significant difference by feed type in both stages (p>0,05) (**Fig. 1-A).** We noticed in the follow-up of flocks that ColR-Kp occurrence in chicken faeces maintained (n=5 flocks) or increased (n=4 flocks) from the P1 to P2 stages, with no significant difference in the ColR-Kp levels between both stages (p>0,05) (**Fig. 1-B**). ColR-Kp was only detected in one of the three chicken meat batches contaminated with *K. pneumoniae* obtained from a farm which was also positive at P1+P2. Regarding environmental samples, ColR-Kp was found in one water sample from one farm, also positive at P1+P2. All isolates were negative for the screening of *mcr* genes but presented MICs for colistin ranging between 4 and ≥16 mg/L, which is compatible with the presence of other acquired resistance mechanisms (results in section 3.5).

### Diversity of *K. pneumoniae* and colistin-resistant *K. pneumoniae*

We recovered 100 1. *K. pneumoniae* (n=50 from OTMF and n=50 from ITMF flocks), from which 56 were colistin-resistant, corresponding to 35 and 20 positive samples, respectively. Most isolates (90%) were recovered from P1 and P2 faecal samples (16 flocks, all farms) (**Fig. 2**). *K. pneumoniae* isolates were grouped into 26 K-types, with twelve dispersed in more than one farm (**Fig. 2; Table S1**). Two K-types represented 40% (n=42) of isolates, independently of feed and included colistin-resistant isolates, KL109 widely dispersed (n=24; 4 farms; P1, P2 and P3; farm B-flock 4 and flock 11 over 4 months) contrasting with KL106 only present in one farm (n=18, water, P1 and P2) (**Fig. 2**). Although less frequent, other K-types were shared by different farms or stages: KL12 (2 farms; P1), KL19 (2 farms; P1, P2 and P3), KL64 (4 farms; P1 and P2), KL111 (2 farms; feed, P1 and P3), KL27 (2 farms, feed and P1) and KL146 (2 farms, P2) (**Fig. 2; Table S1**). A higher diversity was observed in the colistin-susceptible isolates (n=44; 22 K-types) than in colistin-resistant isolates (n=56; 13-Kypes), with 9 K-types shared in both groups.

**Fig. 2.**
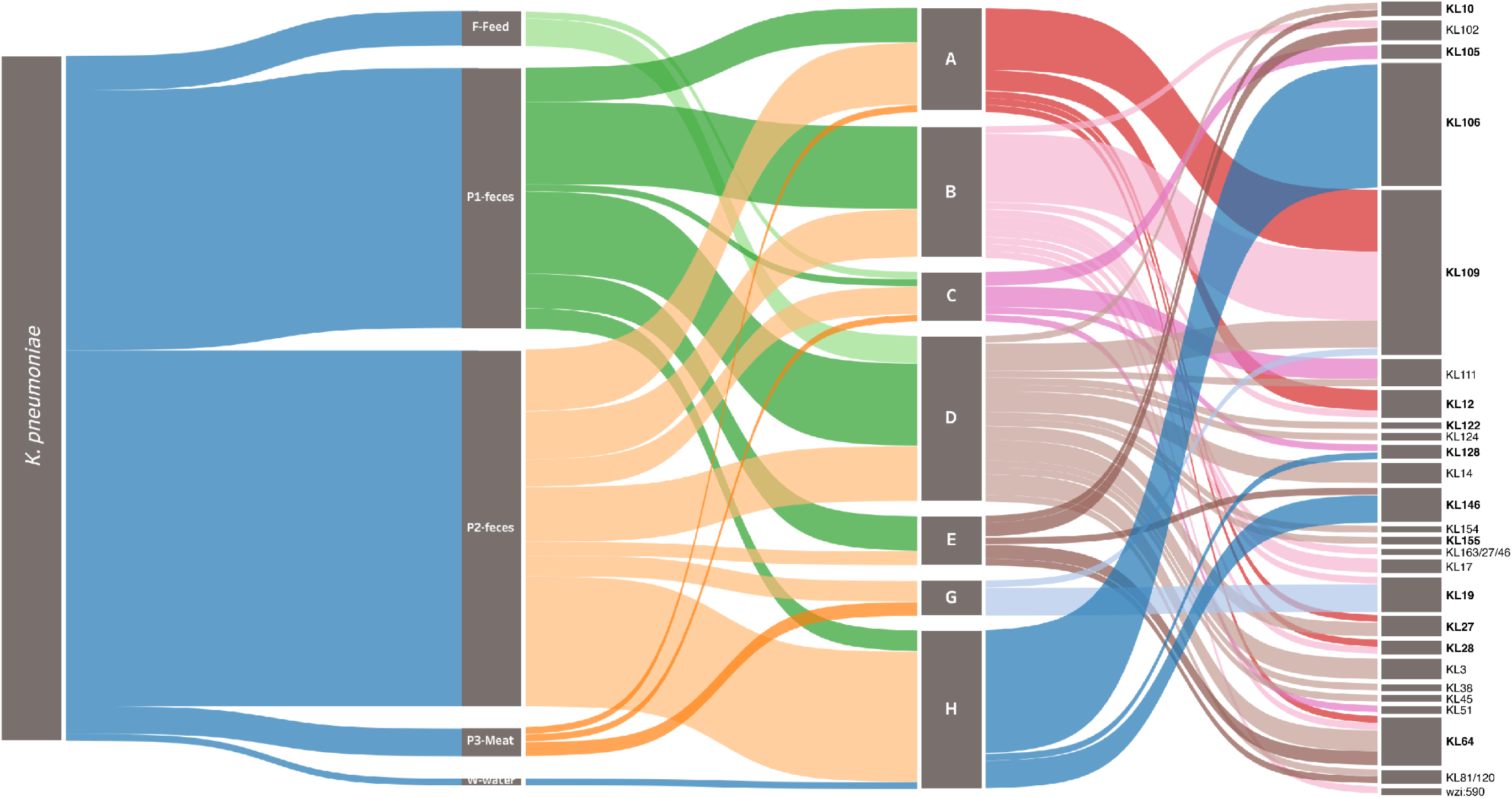
Sankey diagram representing, from left to the right, the occurrence and diversity of *K. pneumoniae* by sample, farm, and K-type. The width of each connection is proportional to the number of positive hits. In bold are indicated the K-types comprising colistin-resistant isolates. The Sankey diagram was generated using Tableau Desktop 2021.4 (https://www.tableau.com/).

### Antibiotic susceptibility of *K. pneumoniae* recovered from poultry production

Considering poultry and meat samples with *K. pneumoniae* (n=31) antibiotic-resistant isolates were observed in all but one sample. Most of these (90%-n=28/31) carried MDR isolates, corresponding to all farms and all but two flocks. Similar MDR and antibiotic resistance rates per sample were observed between both types of feed (p>0,05) (**Fig. 3**). More than 50% of the samples presented at least one *K. pneumoniae* with decreased susceptibility to ciprofloxacin (90%-n=28/31) or resistance to tetracycline, sulphonamides, or trimethoprim (87%-n=27/31, each), gentamicin (65%-n=20/31), colistin (61%-n=19/31) or chloramphenicol (55%-n=17/31) (**Fig. 3).** Regarding the environmental samples with *K. pneumoniae* [feed (n=5; 3 samples) and water (n=1; 1 sample)] all carried MDR isolates. More than 15% of samples (including faeces, chicken meat and water) from diverse farms/flocks presented at least one isolate resistant to extended-spectrum-cephalosporins.

**Fig. 3.**
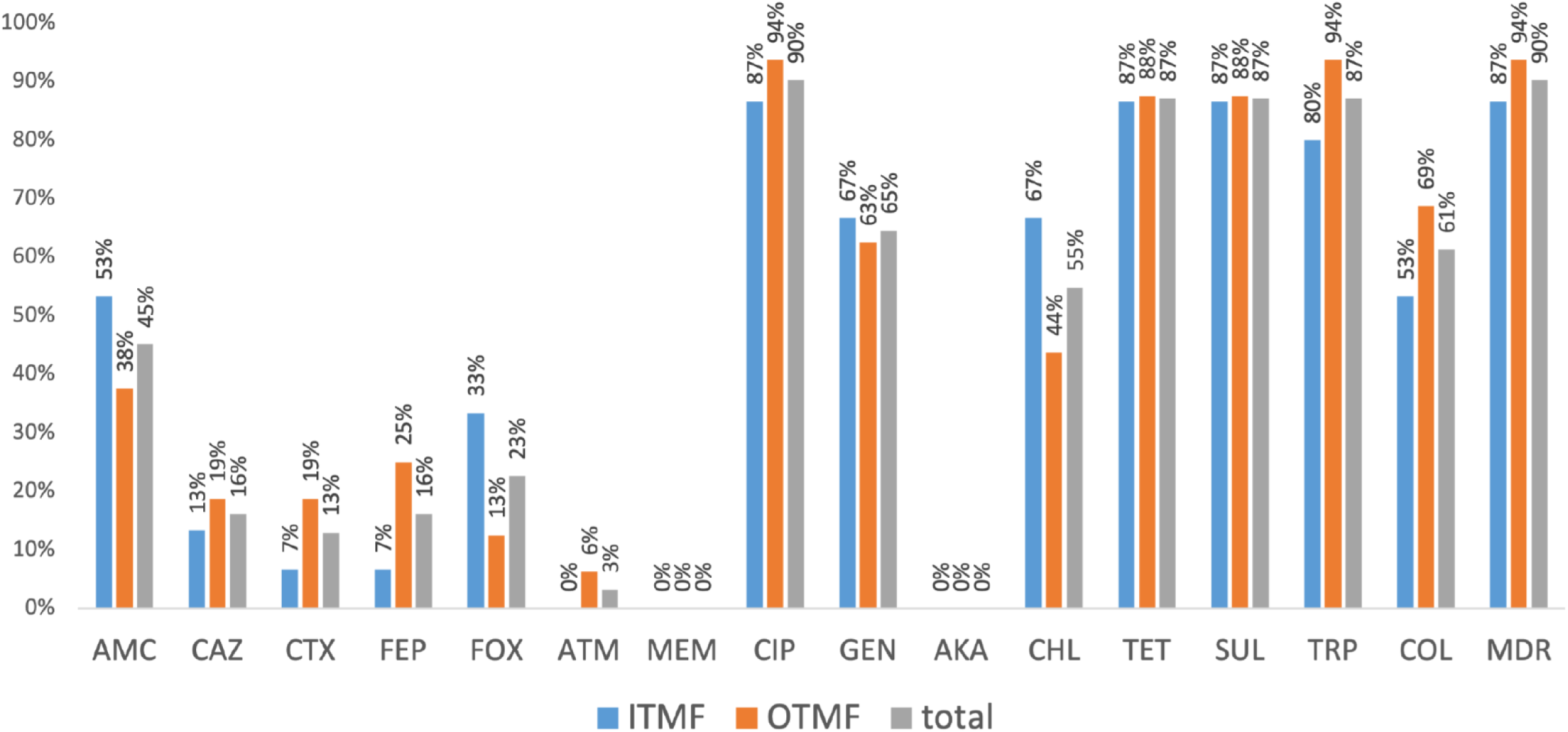
Occurrence of antibiotic-resistant *K. pneumoniae* among positive poultry samples (P1, P2 and P3) by type of feed (ITMF or OTMF). P>0,05 (Fisher exact test). Abbreviations: AMC, amoxicillin+clavulanic acid; CAZ, ceftazidime; CTX, cefotaxime; FEP, cefepime; FOX, cefoxitin; ATM, aztreonam; MEM, meropenem; CIP, ciprofloxacin; GEN, gentamicin; AKA, amikacin; CHL, chloramphenicol; TET, tetracycline; SUL, sulphonamides; TRP trimethoprim; COL, colistin; MDR, multidrug resistance.

*K. pneumoniae* isolates resistant to colistin (n=56/100) were mainly recovered in SCAi media supplemented with this antibiotic, with or without the previous enrichment step (n=30 and n=24, respectively) (**Table 1**). High resistance rates were observed to ciprofloxacin (86%), tetracycline (79%), sulphonamides (74%), or trimethoprim (75%) and MDR (80%), independently of the selection strategy or colistin phenotype (**Table 1**). The MDR isolates belonged to different K-types (n=23), being the most frequent phenotype to ciprofloxacin, tetracycline, sulphonamides, and trimethoprim (n=62 isolates/62%; 18 K-types), associated or not with colistin (n=33 isolates; 9 K-types) and/or extended-spectrum cephalosporins (26 isolates; 7 K-types) resistance (Table S1).

**Table 1.** Antimicrobial susceptibility of Klebsiella pneumoniae (n=100), including colistin-resistant K. pneumoniae (n=56) recovered from poultry production samples

| Isolate selection | *K. pneumoniae*<br>(no. isolates;<br>no. samples) | Antimicrobial resistance (no. of isolates; %)<sup>a</sup> |  |  |  |  |  |  |  |  |  |  |  |  |  |  |  |  |
| --- | --- | --- | --- | --- | --- | --- | --- | --- | --- | --- | --- | --- | --- | --- | --- | --- | --- | --- |
|  |  | COL | AMC | CAZ | CTX | FEP | FOX | ATM | MEM | CIP | AKA | GEN | CHL | TET | SUL | TRP | MDR | pco+sil<br>+ |
| SCAi – Without enrichment/direct | COL-R (n=1; 1) | 1 | 1 | 1 | 0 | 0 | 1 | 0 | 0 | 1 | 0 | 1 | 1 | 0 | 1 | 1 | 1 | 2 |
|  | COL-S (n=6; 5) | 0 | 5 | 0 | 0 | 0 | 1 | 0 | 0 | 6 | 0 | 2 | 4 | 6 | 6 | 6 | 6 | 5 |
|  | **Total (n=7; 6)** | **1 (14%)** | **6 (86%)** | **1 (14%)** | **0** | **0** | **2 (29%)** | **0** | **0** | **7 (100%)** | **0** | **3 (43%)** | **5 (71%)** | **6 (86%)** | **7 (100%)** | **7 (100%)** | **7 (100%)** | **7 (100%)** |
| SCAi – With enrichment/BPW | COL-R (n=1; 1) | 1 | 0 | 0 | 0 | 0 | 0 | 0 | 0 | 1 | 0 | 0 | 0 | 1 | 0 | 0 | 0 | 0 |
|  | COL-S (n=3; 3) | 0 | 0 | 0 | 0 | 0 | 0 | 0 | 0 | 3 | 0 | 1 | 2 | 3 | 3 | 3 | 3 | 3 |
|  | **Total (n=4; 4)** | **1 (25%)** | **0** | **0** | **0** | **0** | **0** | **0** | **0** | **4 (100%)** | **0** | **1 (25%)** | **2 (50%)** | **4 (100%)** | **3 (75%)** | **3 (75%)** | **3 (75%)** | **3 (75%)** |
| SCAi+COL (3.5 µg/mL) - Without enrichment/direct | COL-R (n=30; 13) | 30 | 13 | 9 | 0 | 2 | 13 | 0 | 0 | 28 | 0 | 15 | 14 | 23 | 22 | 21 | 24 | 16 |
|  | COL-S (n=21; 18) | 0 | 3 | 0 | 1 | 1 | 2 | 0 | 0 | 16 | 0 | 5 | 8 | 14 | 11 | 12 | 15 | 17 |
|  | **Total (n=51; 24)** | **30 (59%)** | **16 (31%)** | **9 (18%)** | **1 (2%)** | **3 (6%)** | **15 (29%)** | **0** | **0** | **44 (86%)** | **0** | **20 (39%)** | **22 (43%)** | **37 (73%)** | **33 (65%)** | **33 (65%)** | **39 (76%)** | **33 (65%)** |
| SCAi+COL (3.5 µg/mL) - With enrichment/BPW | COL-R (n=24; 14) | 24 | 10 | 7 | 2 | 3 | 6 | 0 | 0 | 20 | 0 | 16 | 10 | 21 | 21 | 21 | 21 | 13 |
|  | COL-S (n=14; 13) | 0 | 3 | 2 | 2 | 1 | 1 | 1 | 0 | 11 | 0 | 4 | 3 | 11 | 10 | 11 | 10 | 11 |
|  | **Total (n=38; 21)** | **24 (63%)** | **13 (34%)** | **9 (24%)** | **4 (11%)** | **4 (11%)** | **7 (18%)** | **1 (3%)** | **0** | **31 (82%)** | **0** | **20 (53%)** | **13 (34%)** | **32 (84%)** | **31 (82%)** | **32 (84%)** | **31 (82%)** | **24 (63%)** |
| **All isolates** | **Total (n=100; 35)** | **56 (56%)** | **35 (35%)** | **19 (19%)** | **5 (5%)** | **7 (7%)** | **24 (24%)** | **1 (1%)** | **0 (0%)** | **86 (86%)** | **0 (0%)** | **44 (44%)** | **42 (42%)** | **79 (79%)** | **74 (74%)** | **75 (75%)** | **80 (80%)** | **67 (67%)** |

<sup>a</sup> Abbreviations: COL, colistin; COL-R, colistin-resistant *K. pneumoniae*; COL-S, colistin-susceptible *K. pneumoniae*; AMC, amoxicillin+clavulanic acid; CAZ, ceftazidime; CTX, cefotaxime; FEP, cefepime; FOX, ceftoxitin; ATM, aztreonam; MEM, meropenem; CIP, ciprofloxacin; AKA, amikacin; GEN, gentamicin; CHL, chloramphenicol; TET, tetracycline; SUL, sulphonamides; TRP trimethoprim; MDR, multidrug-resistance; SCAi, Simons-Citrate agar with 1% inositol, BPW, Buffered Peptone Water.

### Copper tolerance of *K. pneumoniae* recovered from poultry production

The copper tolerance genes *silA+pcoD* were observed in isolates dispersed by all farms, in most flocks (88%, 14/16) and poultry samples with *K. pneumoniae* (81%, n=25/31) (**Fig. 1-A).** Similar rates were observed between ITMF and OTMF samples among different sampling periods (p>0,05). The *silA+pcoD* were detected in 67% (n=67/100) of the *K. pneumoniae* isolates, regardless of their origin by feed source (n=35/50-70%-ITMF vs n=32/50-64%-OTMF), or farm stage (n=28-P1 vs n=30-P2). The *silA+pcoD*-positive isolates were also detected in chicken meat (n=3; 2 samples-P3), environmental samples [feed (n=5; 3 samples), and water (n=1; 1 sample)]. The *silA*-*pcoD*-positive isolates belonged to 25 out of 26 K-types compared to the *silA-pcoD*-negative that only belonged to 9 K-types. Regarding antibiotic susceptibility, MDR was mostly found in *silA-pcoD*-positive *K. pneumoniae* (91%-n=61/67 *versus* 58%-n=19/33 without *silA+pcoD*; p<0,05), with those, also more resistant to amoxicillin-clavulanic acid, ceftazidime (including ESBL-producers), chloramphenicol, tetracycline, or trimethoprim (p<0,05) (**Fig. 4**). The correlation between colistin resistance and *silA-pcoD*-negative is explained by the high frequency of the colistin-resistant K-type KL109, which corresponded to 23 out of 33 *silA-*negative isolates (**Fig. 4, Table S1**).

**Fig. 4.**
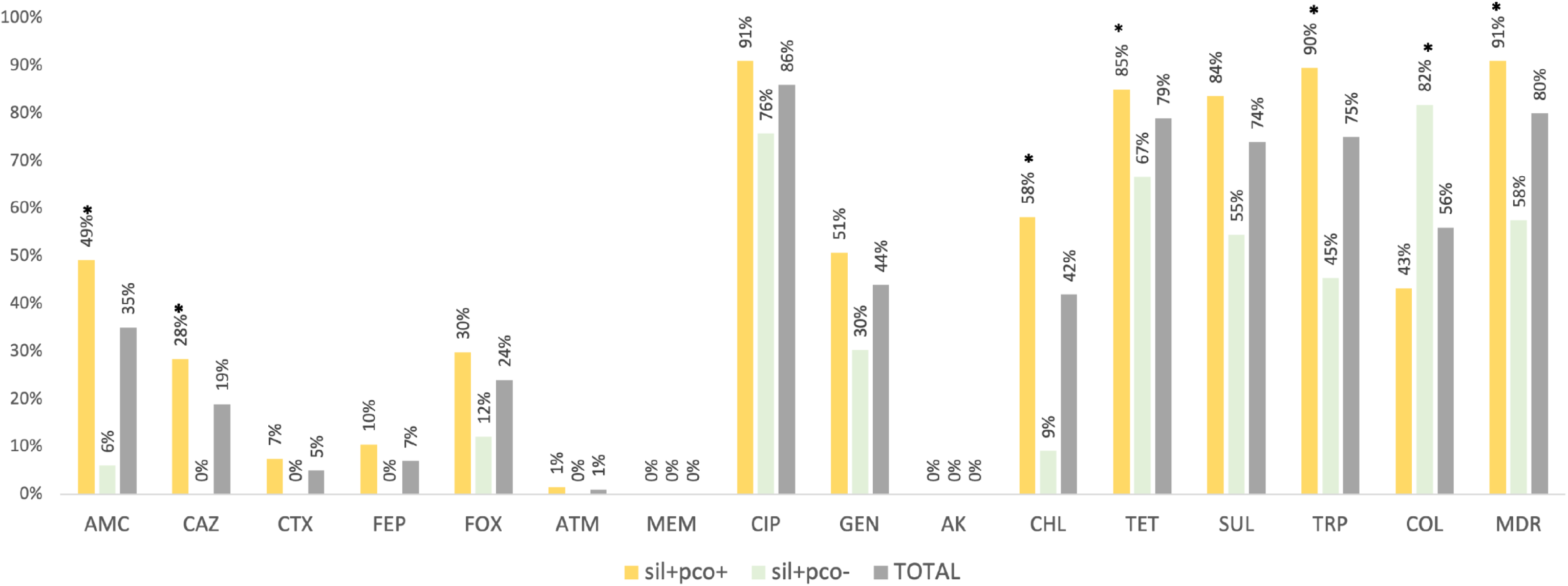
Percentage of antibiotic resistance detected among *pco+silA*+ (n=67) and *pco+silA*-(n=33) *K. pneumoniae* isolates. *p<0,05 (Fisher exact test). Abbreviations: AMC, amoxicillin+clavulanic acid; CAZ, ceftazidime; CTX, cefotaxime; FEP, cefepime; FOX, cefoxitin; ATM, aztreonam; MEM, meropenem; CIP, ciprofloxacin; GEN, gentamicin; AKA, amikacin; CHL, chloramphenicol; TET, tetracycline; SUL, sulphonamides; TRP trimethoprim; COL, colistin; MDR, multidrug resistance.

Copper phenotypic assays were performed in 85% of *K. pneumoniae* carrying or not copper tolerance genes representative of different farms, flocks, K-types, and antibiotic resistance profiles (**Table 2**). All *K. pneumoniae* carrying the *silA-pcoD* genes (100%-n=53/53) exhibited MICs to CuSO_4_ between 16-32 mM (CuT phenotype MIC_CuSO4_ ≥16mM). These data contrast with that of most *K. pneumoniae* without acquired copper tolerance *silA-pcoD* genes (n=30/32), showing MIC to CuSO_4_ of 2-12mM (**Table 2**). The MIC_CuSO4_ distribution was similar for isolates of both feed sources (ITMF *versus* OTMF).

**Table 2.** CuSO_4_ minimum inhibitory concentrations in anaerobiosis for Klebsiella pneumoniae isolates (n=85) by poultry feed type

| Feed | *sil+pco*<br>genes | No. isolates<sup>a</sup> | MIC (mM)<sup>b</sup> |  |  |  |  |  |  |  |  |  |  |  |
| --- | --- | --- | --- | --- | --- | --- | --- | --- | --- | --- | --- | --- | --- | --- |
|  |  |  | 0.5 | 1 | 2 | 4 | 8 | 12 | 16 | 20 | 24 | 28 | 32 | 36 |
| ITMF | + | 29 | - | - | - | - | - | - | 7 | 7 | 12 | 3 | - | - |
|  | - | 14 | - | - | - | 1 | 4 | 8 | 1 | - | - | - | - | - |
| OTMF | + | 24 | - | - | - | - | - | - | 4 | 6 | 12 | 1 | 1 | - |
|  | - | 18 | - | - | 1 | 5 | 1 | 10 | 1 | - | - | - | - | - |
| **Total** | + | **53** | - | - | - | - | - | - | **11** | **13** | **24** | **4** | **1** | - |
|  | - | **32** | - | - | **1** | **6** | **5** | **18** | **2** | - | - | - | - | - |

<sup>a</sup>, The 85 isolates were selected to represent diverse farms, flocks, feeds, copper genotypes and genomic backgrounds.
<sup>b</sup>, *E. coli* ED8739 (plasmid pRJ1004 with *sil+pco* genes; MIC<sub>CuSO<sub>4</sub></sub> 16-20mM, anaerobiosis) and *E. faecium* BM4105RF (negative for all genes tested; MIC<sub>CuSO<sub>4</sub></sub> 2-4 mM, anaerobiosis) were used as control strains in Cu assays (Mourão et al, 2016). Corresponding CuSO<sub>4</sub> mg/L values for the mM values tested are: 0.5 mM, 79.80 mg/L; 1 mM, 159.61 mg/L; 2 mM, 319.22 mg/L; 4 mM, 638.44 mg/L; 8 mM, 1276.87 mg/L; 12 mM, 1915.308 mg/L; 16 mM, 2553.74 mg/L; 20 mM, 3192.18 mg/L; 24 mM, 3830.62 mg/L; 28 mM, 4469.05 mg/L; 32 mM, 5107.49 mg/L; and 36 mM, 5745.92 mg/L.

### Whole-genome analysis of *K. pneumoniae* poultry-associated isolates

The 20 sequenced *K. pneumoniae* representing the most abundant K-types were assigned to 12 STs (10 known and 2 new ST) and 12 lineages based on core genome and KL-types including the globally dispersed ST11-KL105, ST15-KL19, ST147-KL64 and ST307-KL102. Based on the cgMLST analysis and the proposed thresholds (<10 allele differences) (30–32), *K. pneumoniae* were clonally diverse, with ten isolates distributed in five clusters (differing by 0-11 SNPs) (**Fig. 5, Table S2**). Those clusters included isolates from different sample types from the same farm: faeces of poultry and derived meat (n=2 ST6406/farm C and n=2 ST6405/farm A) as well as water and poultry faeces (n=2 ST11/farm H). Also, we detected clones persisting over time in different farms (n=2 ST15/farms E and H; n=2 ST631/farms B and D). The phylogenetic relationship of our genomes with others from different sources, regions, and timeframes available at Pathogenwatch was explored (**Fig. 6**) revealing less than 10 alleles differences in all but four of the lineages (ST147, ST525, ST631, ST2039-2LV). Of remark, isolates belonging to ST15-KL19, ST15-KL146, ST392-KL27 and ST1537-KL64 lineages revealed <21 SNPs with genomes from diverse origins (**Fig. 6**) (a threshold recently proposed for *K. pneumoniae* transmission in healthcare settings) (33). ST15 isolates were linked to genomes associated with human infections in the UK (ST15-KL19), human infections and horses in Italy, France, and the USA (ST15-KL146). ST392-KL27 isolate was linked to genomes from human infections and/or colonization in Spain, while ST1537-KL64 was detected in food products (chicken meat and salads) in France (**Fig. 6**).

**Fig. 5.**
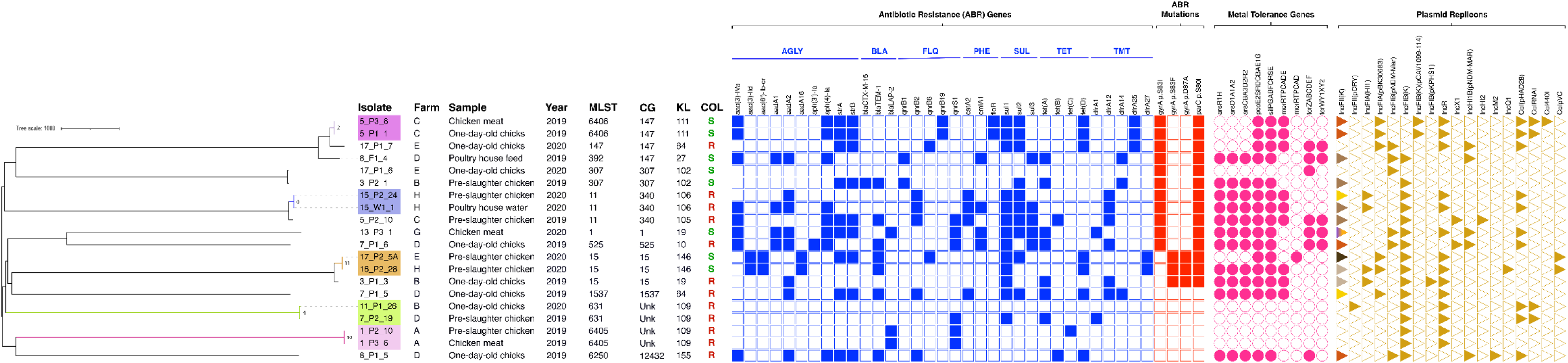
Neighbour-joining tree representing the phylogenetic relationships among the 20 *K. pneumoniae* genomes. The tree was constructed from the Pathogenwatch pairwise-distance matrix (i.e., based on single nucleotide polymorphisms-SNPs called in 1972 core genes). Scale bar units represent substitutions per variant site. The number of substitutions between our isolates from the 5 main clusters are represented in each branch. Associated metadata of all isolates was added using iTOL (https://itol.embl.de/). Each coloured-filled shape represents the presence of relevant antibiotic resistance, metal-tolerance genes, and plasmid replicons associated with well-defined incompatibility groups. The different shades of colours represent the typing results for IncFII(K) plasmids (orange-IncFII_K2_, brown-IncFII_K4_, beige-IncFII_K5_, dark brown-IncFII_K7_, half violet/orange-IncFII_K8_, yellow-IncFII_K21-like_). Only known mutations conferring fluoroquinolone resistance are presented. *Klebsiella* intrinsic antibiotic resistance (*bla*_SHV-1_, *bla*_SHV-11_, *bla*_SHV-26_, *bla*_SHV-28_, *fosA*, *oxqAB*) and metal tolerance (*arsBCR*, *cusABFCRS*) genes were not represented. Abbreviations: AGLY, aminoglycosides; BLA, β-lactams; CG, Clonal Group; COL, colistin; FLQ, fluoroquinolones; KL, K-Locus; MLST, Multilocus Sequence Typing; PHE, phenicols; SUL, sulphonamides; TET, tetracycline; TMT, trimethoprim; Unk, unknown.

**Fig. 6.**
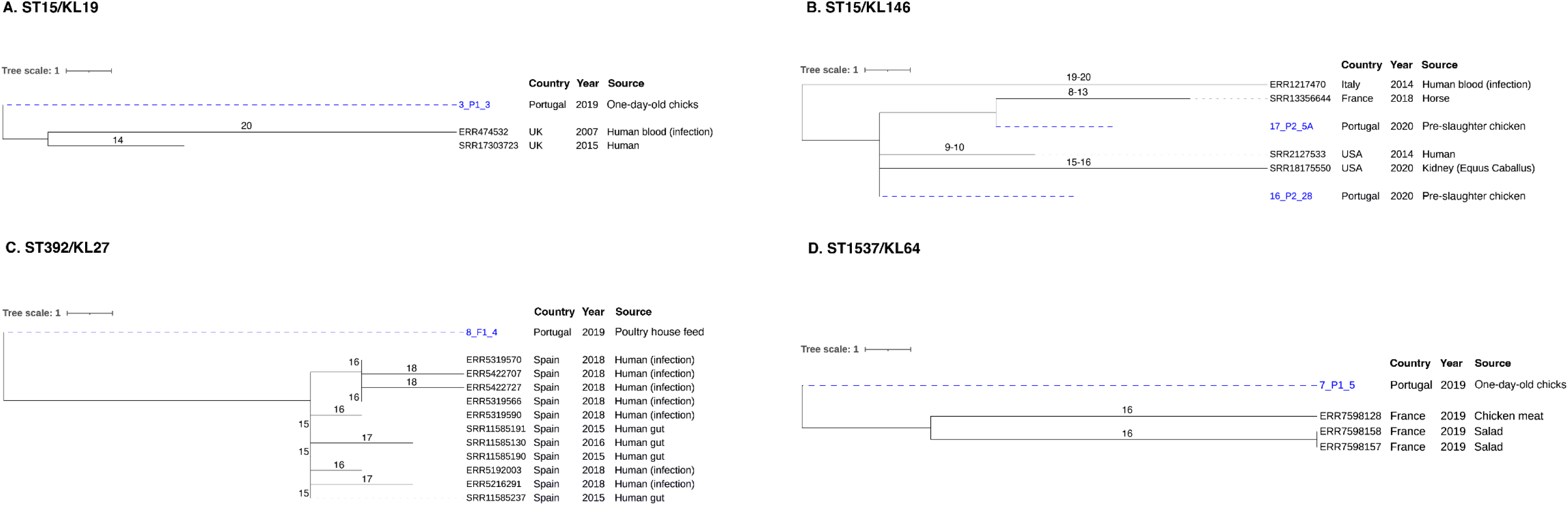
Neighbour-joining trees representing the phylogenetic relationships among our *K. pneumoniae* genomes and those available in Pathogenwatch with less than 21 SNPs. **(A)** ST15/KL19. **(B)** ST15/KL146. **(C)** ST392/KL27. **(D)** ST1537/KL64. The genome selection was performed using the cgMLST single linkage clustering to include the ones with less than 10 allele differences (threshold=10). Then these genomes were used to infer a neighbour-joining tree from the Pathogenwatch pairwise-distance matrix (i.e., based on single nucleotide polymorphisms-SNPs called in 1972 core genes). Scale bar units represent substitutions per variant site. The number of substitutions between our isolates and the ones available in Pathogenwatch are represented in each branch. All isolates’ associated metadata (country, source of isolation and collection date) was added using iTOL (https://itol.embl.de/).

The WGS revealed a high load and diversity in antibiotic resistance and metal tolerance compared to virulence genes (**Fig. 5, Table S3**). All isolates carried the chromosomally encoded *mrkABCDFHIJ* cluster encoding for a type 3 fimbriae, whereas only ST15 isolates carried the virulence accessory genes *kfuABC* (ferric uptake system) and the *kpiABCDEFG* (pili system). Regarding the genomic analysis of colistin resistance, we detected 96 different chromosomal mutations (mostly missense; n=94/96) in 72% of the genes (n=23/32) encoding for proteins previously implicated in colistin resistance compared with the reference strain *K. pneumoniae* MGH 78578 (described in detail in **Fig. 7** and **Fig. S3**). From these, 52 distinct mutations were present in all ColR-Kp (9 lineages) across five operons/genes associated with colistin resistance mechanisms such as modifications of LPS/lipid A (*arnABDFT, crrAC, phoQ, pmrACD, mgrB*), overexpression of efflux pumps (*acrB*), LPS loss (*lpxA, msbA*) or biosynthesis (*yciM*), and regulation (*rstB*) (**Fig. 7**). Diverse mutations implicated in colistin resistance were detected in seven of those genes (*acrB*, *arnB*, *arnD*, *arnT*, *mgrB*, *phoQ* and *pmrC*), varying accordingly with the clone (**Fig. 7**). All but one of the isolates accumulated mutations in genes from different clusters (varying from two to seven).

**Fig. 7.**
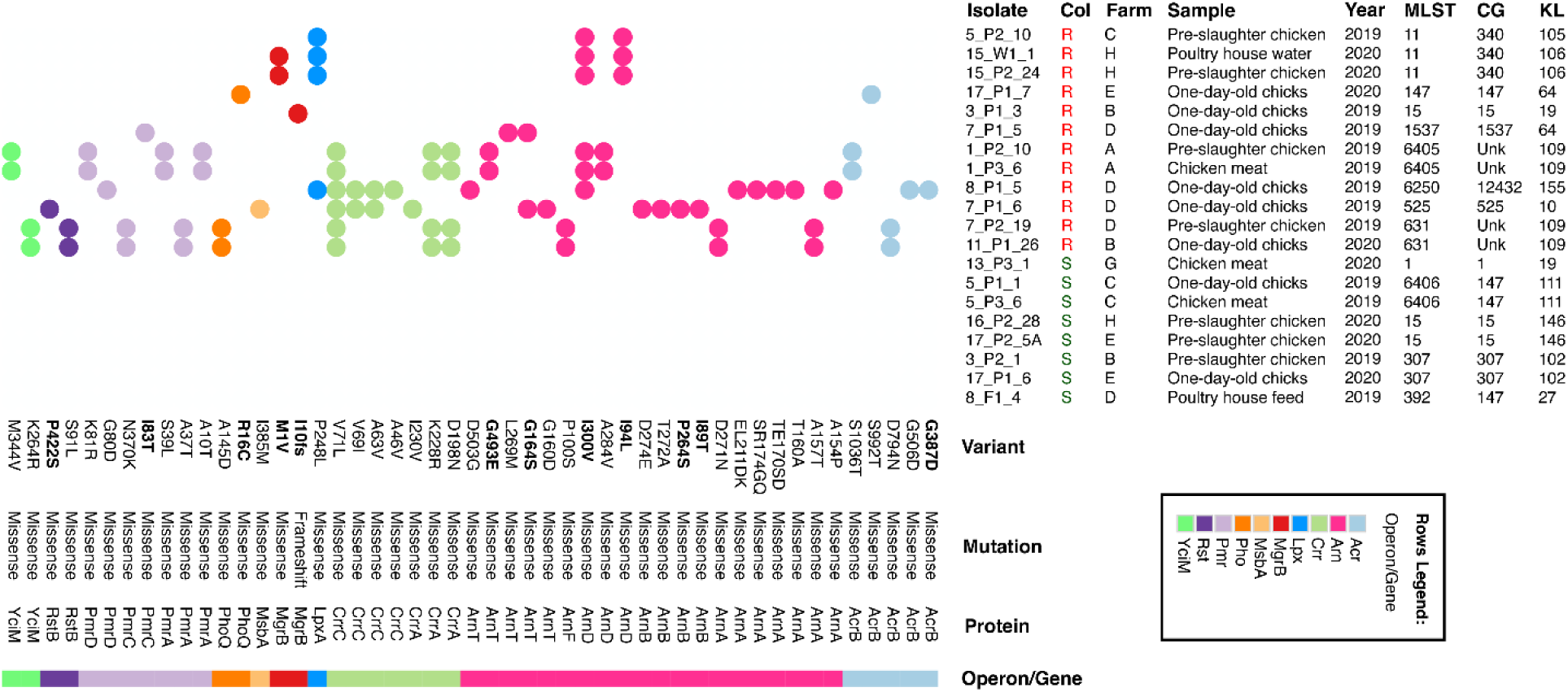
Heatmap generated using Morpheus online web-server (https://software.broadinstitute.org/morpheus/) representing the distribution of distinct mutations among the sequenced colistin-resistant *Klebsiella pneumoniae* genomes. Coloured squares indicate the presence of a specific mutation, while each colour represents the genes that are part of the same operon. Variants in bold were supported by the available literature as present in ColR-Kp and/or were predicted as deleterious by PROVEAN. *Klebsiella pneumoniae* isolates were grouped into colistin-susceptible when MIC = <2 μg/mL or colistin-resistant when MIC = ≥2 μg/mL. When required, figures were minimally edited manually using Adobe Illustrator v25.3.1. Detailed resistance mechanisms: *acrB* – efflux pump; *arnABDFT* - lipid A modification with L-Ara4N addition; *crrA* – lipid A modification by upregulation of *pmrAB*/activation of the glycosyltransferase; *crrC –* connector protein; *lpxA* – inactivation of lipid A biosynthesis abolishing LPS synthesis; *mgrB* - inactivation of negative feedback regulator of the PhoP/PhoQ system; *msbA* - ABC transporter of lipid A; *phoQ* and *pmrA* - activation of LPS-modifying operation in the two-component systems; *pmrC* - lipid A modification with PEtn; *pmrD* – connector protein; *rstB* - sensor protein; *yciM* - regulation of LPS biosynthesis. Abbreviations: A, alanine; C, cysteine; D, aspartate; E, glutamate; G, glycine; I, isoleucine; K, lysine; L, leucine; M, methionine (start codon); N, asparagine; P, proline; R, arginine; S, serine; T, threonine; V, valine; Unk, unknown.

Beyond colistin resistance, we detect diverse acquired genes (n=36) encoding resistance to seven different antibiotic classes, with 85% of genomes (n=16/20) carrying ≥4 classes but varying between lineages (0-6). The most frequent classes and genes were aminoglycosides [*strA*/*strB*], sulfonamides (*sul1/sul2*), tetracyclines [*tet(A)*/*tet(D)*], trimethoprim (*dfrA*) and phenicols (*catA, cmlA*) (**Fig. 5, Table S3**). Also, analysis of the genetic background of resistance to ciprofloxacin showed chromosomal mutations in the quinolone resistance determining region (QRDR) of topoisomerase genes *gyrA* (S83I, S87A and S83F) and *parC* (S80I) (*gyrA*-*parC*, 70%-n=14/20) and variable plasmid-mediated quinolone resistance (PMQR) *qnrB* (n=11) and *qnrS* (n=6) genes (**Fig. 5**). Extended-spectrum cephalosporin *bla*_CTX-M-15_ was detected in only one ST307 isolate. Moreover, diverse acquired metal tolerance gene clusters encoding for copper/silver (*pco/sil*), arsenic (*ars*), mercury (*mer*) and/or tellurite (*ter*) tolerance were detected in all but four of the genomes (**Fig. 5**). The *pco*+*sil* cluster (75%-n=15/20) was frequently associated with operons *ars* (73%-n=11/15), *mer* (67%-n=10/15) and/or *ter* (40%-n=6/15) (**Fig. 5**). Co-occurrence of antibiotic resistance genes and copper tolerance genes in 75% (n=15/20) of the genomes was detected. Genomes carried an average of 5 plasmids (1-9 per genome), and a total of eighteen different well-defined plasmid incompatibility groups (1-7 replicon types per isolate). The most common were IncFIB_K_ (80%; n=16/20), followed by diverse IncFII_K_ (70%; n=14/20) and IncR (70%; n=14/20), most often in variable combinations with FIA, other FIB types (FIB_pNDM-MAR_, FIB_pCAV1099_, FIB_pKPHS1_) and/or HI1B (**Fig. 5**). Col plasmids were also frequent (65%; n=12/20 genomes) while other classical incompatibility groups were rare (IncHI2, IncX, IncQ, IncM). Copper tolerance genes *pco*+*sil* were located mainly in IncFIB_K_+IncFII_K_ plasmids (73%; n=11/15, ∼80-270 Kb) of different IncFII_K_ groups by pMLST (2, 4, 5, 7, 8, 21-like) alongside with variable antibiotic resistance genes (*aac(3)-IVa*, *aph(4)-Ia*, *aadA2*, *strA*-*strB*, *qnrB1*, *catA2*, *sul1*, *sul2*, *tetA*, *tetD*, *dfrA12*, *dfrA14*, *bla*_TEM_, *bla*_CTX-M-15_) (**Fig. 5**). These plasmids carrying *pco*+*sil* were similar to others (MOB-Recon, mash distance 0.0012 – 0.0309) described in humans in multiple countries, occasionally carrying *bla*_CTX-M-15_, *bla*_DHA-1_ and/or *qnr* (**Table S4**).

## DISCUSSION

This study firstly showed the absence of *mcr* but a high occurrence and diversity of colistin-resistant (*mcr*-negative) *K. pneumoniae* after >2 years of colistin withdrawal in intensive chicken farms, independent of the type of Cu-supplementation used in the feed formulation. Furthermore, we demonstrate the persistence of particular *K. pneumoniae* clones throughout the whole poultry production chain and suggest poultry as a potential foodborne/environmental source of *K. pneumoniae* lineages with clinical relevance.

The high rates of intensely raised flocks positive for *K. pneumoniae* in the early and pre-slaughter stages, together with the detection of *K. pneumoniae* with or without standard faecal indicators in feed or water samples (e.g., *E. coli* and/or *Enterococcus* detected in 9 out of 14 water samples from all farms) (data not shown), suggests diverse contamination events (e.g., hatchery farm, poultry-house cleanliness and biosecurity level, water, feed, inanimate surfaces and/or human handlers) occurring early and frequently along the production chain (34, 35). Environmental factors (e.g., type of litter, temperature, relative humidity, daily cycles, moment of sampling/age, season, vacancy period) or the use of antimicrobial compounds (e.g., disinfectants, antibiotics, coccidiostats or metals) are also known to have an impact on the composition of poultry gut microbiota (35–37). These factors could justify the differences found in faecal *K. pneumoniae* occurrence rates at the farm level in this study (82%, varying from 25-100%) and compared to another recent study using also SCAI media for bacterial recovery in chicken farms (26%) (38). However, despite the high rate of faecal samples carrying *K. pneumoniae*, resistant or not to colistin, our data revealed few positive chicken meat batches (17%), which is far below what was reported in studies from the USA (47%) and the EU (60%) (39, 40). Besides, *Salmonella* was not detected in any sample (data not shown). These results demonstrate reduced meat cross-contamination and the effectiveness of sanitary measures during animal transport and at the slaughterhouse, thus reducing consumer’s risk of exposure (12, 37, 41).

Recent studies suggest that banning colistin in food-animal production has had an encouraging outcome by limiting *mcr* spread (11, 12). Our results confirm this trend in poultry production by the absence of *mcr* in farms and meat isolates, although a high rate of samples carried ColR-Kp associated with a high diversity of chromosomal mutations, independently of the feed type used. This suggests the circulation of diverse colistin-resistant genotypes responding differently to the colistin ban. We detected mutations, alone or in combination, in genes known to be involved in lipid A modifications and colistin resistance phenotypes (*arnABDFT, phoQ, pmrACD* and *mgrB*), as described (42–50). However, a high variety of other non-described mutations were detected, supporting the urgent need for reliable genotypic-phenotypic correlations to explain colistin resistance mechanisms (51). Such a variety and frequency of chromosomal mutations in ColR-Kp detected in different studies and environments (26, 43, 50, 51) suggest they could play a role in adaptive features other than colistin resistance (e.g., *mgrB* inactivation in environmental *K. pneumoniae* survival and transmission; *phoPQ* or *pmrB* mutations in chlorhexidine tolerance) (52–54). Also, some of these mutations do not confer significant biological cost (55, 56) which may explain the occurrence of colistin-resistant *mcr*-negative *K. pneumoniae* long after colistin withdrawal. Finally, other factors contributing to co-selection (e.g., other antibiotics, growth-promoters such as Cu) or maintenance (continuous contamination sources) of chromosomally-mediated colistin-resistant strains in the livestock sector (26), could not be discarded.

In contexts of antibiotic/colistin reduction and replacement, the use of diverse antimicrobial compounds may be expanded, making it difficult to understand the complexity of possible co-selection events of the MDR strains. Some recent studies performed after colistin withdrawal revealed an increase in clinically relevant antibiotic resistance genes (e.g., *bla*_CTX-M_ and *bla*_NDM_) among food-animal isolates (57, 58). In this study, these genes were not detected but we show high rates of resistance to multiple antibiotics in *K. pneumoniae* isolates from poultry, independently on the sample selection strategy (with or without colistin supplementation) or between sample groups (poultry life stages and feed type). Resistance rates were higher for antibiotics (e.g., ciprofloxacin, tetracycline) frequently used for therapeutics at poultry farms (12, 59), in contrast with other studies in poultry or other non-clinical environments (38, 60) in which resistance rates were low. Such differences in the available studies could reflect local variation in the usage of several antibiotic classes, the diversity of routes for *K. pneumoniae* dissemination within and beyond the production environment, and the circulating clonal lineages (38, 60).

Other largely used feed ingredients with antimicrobial activity like metals/copper may also contribute to changes in poultry microbiota and/or even potentiate Cu-tolerant and MDR strains co-selection (19, 20, 24), a factor unexplored for *K. pneumoniae*. Recent studies revealed correlations between metal environmental pollution (e.g., Cu at poultry farm samples) and an increase in antibiotic resistance genes (61, 62). We have also detected the plasmid-acquired copper tolerance *pco+sil* cluster in isolates dispersed in all farms and in most of the flocks. Furthermore, we describe for the first time in *K. pneumoniae*, a correlation between the presence of *pco+sil* genes and the Cu tolerance phenotype (MIC≥16mM in anaerobiosis). These data suggest the adaptability of this species to stressful/unfavourable farming environments, as previously detected for other zoonotic bacteria such as the emergent MDR clones of *Salmonella* (63). According to our data, the feeding regime (ITMF or OTMF) does not seem to contribute to the expansion of MDR or ColR-Kp, suggesting also a similar whole farm environment and management practices. Thus, studies evaluating the environmental factors (e.g., pH) (64) and sub-inhibitory concentrations of heavy metals to maintain antibiotic-resistant bacteria in poultry production and other related environments (61, 65, 66) are urgent to clarify the persistence of MDR-*K. pneumoniae*, namely to colistin, carrying heavy metal tolerance genes.

It is of remark that metal tolerance operons (*pco*+*sil*) were mainly located in multireplicon F-type (FII_K_+FIB_K_) plasmids carrying genes encoding resistance to several classes of antibiotics, including the critical extended-spectrum cephalosporins and/or fluoroquinolones, supporting the potential for diverse co-selection events (67, 68). These mosaic F-type plasmids are common in *K. pneumoniae* populations from different sources (68–73), suggesting a major role in both dissemination and persistence of antibiotic resistance and metal tolerance genes, and *K. pneumoniae* adaptation to different niches. Besides, the similarity between plasmid backbones identified in poultry isolates with those described in *K. pneumoniae* collections from humans (**Table S3, Table S4**) (72) suggests a common pool of shared plasmids between humans and eventually different animal species (70) that deserves to be further explored with comprehensive comparative plasmidome analysis.

This is the first study tracking *K. pneumoniae* throughout the whole poultry production chain, enabling the identification of multiple transmission routes in flocks and the farm environment. The identification of closely related MDR *K. pneumoniae* isolates (e.g., ST11-KL106, ST15-KL146, ST631-KL109, ST6405-KL109 and ST6406-KL111) in multiple poultry stages and environments in the same or different farms, suggests that food sector efforts should be placed in improving the sanitary measures in the poultry-houses, poultry-workers, and the environment beyond (treatment of wastewaters and manure). The persistence of MDR *K. pneumoniae* clones in the poultry chain, which is according to previous studies conducted on farms (34, 35, 38) or slaughterhouses (41), suggests the presence of adaptive environmental features other than antimicrobial resistance (e.g. biofilm formation). Most available studies assessing *K. pneumoniae* sources from a One Health perspective have included retail foods, livestock farms or wastewater while the poultry chain has been greatly understudied (27, 38–40, 59, 60, 74, 75). Despite the high diversity found, we highlight the commonality of KL-types (KL64, KL102, KL105, KL106) and lineages (ST15-KL19, ST15-KL146 and ST392-KL27) identified between poultry and human clinical isolates (this study) (28, 76). Although the immunogenicity of different KL-types is not well understood, particular capsular types have been shown to influence the selection of certain *K. pneumoniae* lineages (77), which might provide an advantage regarding host immunity (78, 79). Finally, despite human contamination cannot be discarded, our data suggest poultry as a reservoir and source of globally dispersed and human clinically relevant *K. pneumoniae* lineages with a potential risk for transmission to humans and other ecosystems (animals and the environment).

## CONCLUSIONS

This study constitutes the first comprehensive analysis of *K. pneumoniae* from a poultry farm-to-fork perspective. Our data revealed, independently of the Cu-feed type, a high rate of chicken faecal samples carrying a high diversity of *K. pneumoniae*, MDR, Cu-tolerant and colistin-resistant/*mcr*-negative strains long after colistin withdrawal. Furthermore, we propose that poultry may serve as a reservoir and source of human clinically relevant lineages and mosaic F-type plasmids (with antibiotic resistance and metal tolerance genes), which could pose a potential risk of human exposure through the food chain (e.g., poultry carcasses, occupational exposure) and/or environmental release (e.g., farm or slaughterhouse wastewaters). Therefore, it is crucial to further explore the drivers contributing to the circulation and persistence of MDR *K. pneumoniae* in poultry production. This will help to identify novel mitigation strategies addressing producers’ efforts, consumer concerns, as well as the EU farm-to-fork goals, in the fight against antimicrobial resistance and towards improving food safety and environmental sustainability.

## MATERIAL AND METHODS

### Sampling strategy at the chicken farm and slaughterhouse processing plant

Our pilot study involved seven Portuguese intensive-based chicken farms with similar conventional indoor and floor-raised production systems, being all the key practices and requirements in compliance with EU legislation, as indicated by the operator’s (80). The farm selection was based on those whose grow-out poultry houses had similar conditions and individual feed silos to be possible to divide each flock at arrival (one-day-old chicks) in half, each one under a different feed type. In all farms, colistin was banned since January 2018, while copper was routinely used as an additive in poultry feed. The current inorganic formulation feed (ITMF) was supplemented with inorganic sources of Cu and the organic formulation (OTMF) with organic forms, previously evidenced as capable of maintaining broiler performance under commercial conditions (23). Both mineral supplements were added to the identical commercial starter and grower feeds adapted to different periods of the broileŕs life, with copper concentrations decreasing ≅42% between ITMF and OTMF feeds (both far below the maximum dose of 25 mg/kg of Cu, according to the EU Regulation 2018/1039).

Eighteen chicken flocks (10000-64000 animals from each house; Ross 308 strain) fed with ITMF (n=9) *versus* OTMF (n=9) were sampled at each farm between October 2019 and November 2020 (two farms were sampled twice in different seasons). Chicken faeces (≅50g; total n=34) were collected in each farm from two separated poultry houses (ITMF *versus* OTMF) at 2-3 days of life (P1 stage; n=18 samples) and after 28-30 days (P2 stage; n=16 samples; 2 samples missing during 2020 COVID-19 lockdown), the day before slaughter. Environmental samples, including feed (≅50g; n=18) and water (≅1L; n=14), were also collected inside grow-out houses at each farm at the P1 stage (**Fig. S1**).

Raw chicken meat samples (n=18 batches; recovered after slaughter and air chilling) of the same flocks were collected after slaughter (age 30-35 days; P3 stage) in the poultry-production slaughterhouses, immediately before distribution for retail sale. Each sample was processed as a pool of neck skin from 10 carcasses of the same batch (each batch corresponded to one flock from the same farm slaughtered at the same time) (**Fig. S1**).

All the previous solid and water samples were collected in sterile plastic bags or containers, transported at 4°C, and processed on the same day at the laboratory. Subsequent sample processing was performed by cultural approaches, as described in the following sections.

### Screening of *K. pneumoniae* and colistin-resistant *K. pneumoniae*

*K. pneumoniae* was selectively recovered, directly from the sample and after enrichment, in the Simmons citrate agar plates with 1% inositol (SCAI), a medium that selectively favours the growth of *Klebsiella* in potential *Escherichia coli*-rich samples (**Fig. S2**). A common initial step consisted of weighing 25g of tested solid samples (P1, P2 and P3 stages) or filtering 500 ml of water samples (P1 stage) into 225 mL of Buffered Peptone Water (BPW) supplemented with 3.5 mg/L of colistin. The direct cultural method included spreading an aliquot of 100 µL of the BPW+colistin after 1 hour at room temperature (resuscitation step) on SCAI supplemented or not with colistin (3.5 mg/L). The enrichment approach involved the same procedure, but after a previous incubation of BPW+colistin, at 37°C for 16-18 h. All the SCAI plates were incubated at 37°C for 48h. One up to five colonies of each presumptive morphotype were selected for identification. Isolates’ identification was performed by Matrix-Assisted Laser Desorption-Ionization-Time of Flight Mass Spectrometry (MALDI-TOF MS) (MALDI-TOF VITEK MS, bioMérieux, France) and by PCR for *K. pneumoniae* (81). In all the identified isolates, screening of colistin resistance genes (*mcr-1-5* and *mcr-6-9*) was assessed by two multiplex PCR, as previously reported (82, 83). The Minimum Inhibitory Concentration (MIC) for colistin was determined by the reference broth microdilution (84). An estimation (CFU/g) of colistin-resistant *K. pneumoniae* in the poultry samples directly plated on SCAI+colistin (see above the procedure) was performed after counting typical *Klebsiella* colonies, species identification and MIC_colistin_.

### Phenotypic and genotypic characterization of *K. pneumoniae*

Relatedness between isolates from different samples was inferred by Fourier Transform Infrared (FT-IR) spectroscopy with attenuated total reflectance (ATR) using a Perkin Elmer Spectrum Two instrument. After growth in standardized culture conditions (37°C/18h), a colony was directly deposited on the ATR accessory of the FT-IR instrument and air-dried. Spectra were acquired in standardized conditions (4000-600 cm^-1^, 4cm^-1^ resolution, and 16 scan co-additions) and then compared between each other and with those from an in-house spectral database (allowing identification of 30 K-types from well-characterized international *K. pneumoniae* clones), as previously described (76, 85). FT-IR-based assignments were confirmed using PCR of the *wzi* gene and further sequencing at Eurofins Genomics (https://www.eurofinsgenomics.eu/) to infer the K-type at BIGSdb (http://bigsdb.pasteur.fr/klebsiella/klebsiella.html) (86).

Antibiotic susceptibility profiles were determined by disc diffusion using fourteen antibiotics (amoxicillin+clavulanic acid-30 µg, amikacin-30 µg, aztreonam-30 µg, cefepime-30 µg, cefotaxime-5 µg, cefoxitin-30 µg, ceftazidime-10 µg, chloramphenicol-30 µg, ciprofloxacin-5 µg, gentamicin-10 µg, meropenem-10 µg, sulfamethoxazole-300 µg, tetracycline-30 µg, and trimethoprim-5 µg). The interpretation was performed using the European Committee of Antimicrobial Susceptibility Testing (EUCAST) (87) and, when this was not possible, by the Clinical and Laboratory Standards Institute (CLSI) guidelines (88). *Escherichia coli* ATCC 25922 was used as the control strain. MDR was considered when the isolates were resistant to three or more antibiotics of different families (in addition to ampicillin to which all *K. pneumoniae* are intrinsically resistant).

All the isolates were screened for the *silA*+*pcoD* Cu tolerance genes by PCR, as previously reported (63). Copper susceptibility was studied in representative isolates (representing the diverse farms, flocks, feeds and genomic backgrounds) by the agar dilution method (MIC_CuSO4_-0.5-36 mM; anaerobiosis), as previously published (63).

### Whole-genome sequencing (WGS) and comparative analysis

Representative isolates (n=20) of different farms, sources, timespans, and clones/K-types were selected for whole genome sequencing (WGS). The DNA was extracted with the Wizard Genomic DNA purification kit (Promega Corporation, Madison, WI), with the final concentration measured with Qubit 3.0 Fluorometer (Invitrogen, Thermo Fisher Scientific, USA) and sequenced with Illumina NovaSeq 6000 S4 PE150 XP (Illumina, San Diego, CA) at Eurofins Genomics (https://eurofinsgenomics.eu/). Raw reads quality control was assessed with default parameters using FastQC v0.11.9 (89) and MultiQC v1.12 (90). Good quality raw reads were then *de novo* assembled using SPAdes v3.15.3 (91) with "--isolate" and "--cov-cutoff auto" flag options. We then used QUAST v5.0.2 (92) and CheckM v1.0.18 (93) within KBase (https://www.kbase.us/) to assess the quality and completeness of the assemblies.

The assemblies were annotated using the RAST server (94). Genomes’ assemblies were then uploaded to Pathogenwatch v2.3.1 (https://pathogen.watch/) to determine species, capsular polysaccharide (K) and lipopolysaccharide O locus types and serotypes (29), multi-locus sequence types (MLSTs) (95) and core genome MLSTs (cgMLST). Pathogenwatch uses the calculated pairwise single nucleotide polymorphisms (SNPs) distances between the genomes based on a concatenated alignment of 1972 core genes to infer a neighbour-joining tree for phylogenetic analysis (96). Closely related genomes and the associated metadata (country, source of isolation and collection date) were retrieved from all public genomes available at Pathogenwatch, after cgMLST single linkage clustering and selection of those with less than 10 allele differences (threshold=10). Neighbour-joining trees were edited using iToL (97).

Snippy v3.2-dev (https://github.com/tseemann/snippy) was used to identify substitutions (SNPs) and insertions/deletions (InDels) in genes (n=32) putatively associated with colistin resistance (43, 51, 98), by aligning the raw read data from each isolate against the reference genome *Klebsiella pneumoniae* MGH 78578 (GenBank accession number CP000647). All the gene mutations were further manually confirmed in the assembled genomes using the Geneious Prime® 2023.0.1 software (http://www.geneious.com/). The deleterious effect of detected mutations on protein function was further assessed using the Protein Variation Effect Analyzer – PROVEAN (99) or data from previous publications. We used the standard PROVEAN scores of lower than or equal to -2.5 for a deleterious effect and higher than -2.5 for a neutral effect on protein function.

Kleborate v2.2.0 (100), integrated into Pathogenwatch, was used for the prediction of antibiotic resistance determinants (acquired and chromosomal mutations), and virulence traits (yersiniabactin-YbST, colibactin-CbST, aerobactin-AbST, salmochelin-SmST, and regulators of mucoid phenotype-RmpA, and RmpA2). ABRicate v1.0.1 (https://github.com/tseemann/abricate) with *in-house* databases was used to detect additional *K. pneumoniae* virulence genes from BIGSdb-Pasteur (https://bigsdb.pasteur.fr/klebsiella/), the chaperone-usher pili system (*kpiABCDEFG*) (101), and relevant metal tolerance genes (*cusABFCRS*, *arsBCR*, *arsR1H-arsD1A1A2-arsCBA3D2R2*, *pcoGE1ABCDRSE2*, *silESRCFBAGP*, *merRTPCADE,* and *terZABCDEF-terWY1XY2Y3* operons). AMRFinderPlus v3.10.30 (102) was used for the determination of antibiotic-resistant novel alleles.

In all the 20 WGS-selected isolates, plasmid replicon typing was determined using PlasmidFinder (103, 104) and pMLST v2.0 (104) from the Centre for Genomic and Epidemiology (http://www.genomicepidemiology.org). IncFII_K_ plasmids were further characterized according to (105)) (https://pubmlst.org/organisms/plasmid-mlst). To confirm the location of metal tolerance (MeT) genes and reconstruct putative plasmids based on draft assemblies, we used the MOB-recon tool v3.0.0 from the MOB-suite package (106, 107). The MeT genes were considered part of a specific plasmid when identified by MOB-recon or when located on the same contig as the replicon/incompatibility determinant.

### Statistical analysis

Differences in occurrence, antimicrobial resistance, copper tolerance and bacterial counts among *K. pneumoniae* and flocks fed with ITMF and OTMF as well as among P1, P2 and P3 stages were analysed by Fisher exact and Wilcoxon matched-pairs signed rank tests (α=0.05) using Prism software, version 8.1.1 (GraphPad).

### Data availability

The data for this study have been deposited in the European Nucleotide Archive (ENA) (https://www.ebi.ac.uk/ena) under BioProject accession number PRJEB60418.

## ACKNOWLEDGEMENTS

The authors would like to thank Lina Cavaco, Yang Wang, Alessandra Carattoli and Maria Borowiak for providing the *mcr*-positive controls, the pre-graduation students Beatriz Pereira (FCNAUP), Carina Baptista (FFUP), Eulália Monteiro (FCNAUP) and Sofia Paiva (FCNAUP) for technical support and the staff of the participating farms and slaughterhouses for their kind cooperation. We thank the Institut Pasteur teams for the curation and maintenance of BIGSdb-Pasteur databases at http://bigsdb.pasteur.fr/.

This work was supported by the Applied Molecular Biosciences Unit - UCIBIO which is financed by national funds from FCT - Fundação para a Ciência e a Tecnologia [UIDP/04378/2020 and UIDB/04378/2020], by the Associate Laboratory Institute for Health and Bioeconomy-i4HB [LA/P/0140/2020], by the AgriFood XXI I&D&I project [NORTE-01-0145-FEDER-000041] co-financed by the European Regional Development Fund (ERDF) through NORTE 2020 (Programa Operacional Regional do Norte 2014/2020) and by the University of Porto and SOJA DE PORTUGAL [Grant N° IJUP-Empresas-2019-17]. Joana Mourão and Ângela Novais were supported by national funds through FCT/MCTES in the context of the Scientific Employment Stimulus (2021.03416.CEECIND and 2021.02252.CEECIND, respectively). Marisa Ribeiro-Almeida and Andreia Rebelo were supported by PhD fellowships from FCT (SFRH/BD/146405/2019 and SFRH/BD/137100/2018, respectively). The authors are greatly indebted to all the financing sources.

**Joana Mourão:** Methodology, Software, Formal analysis, Investigation, Visualization, Writing – original draft, Writing – review & editing. **Marisa Ribeiro-Almeida:** Methodology, Formal analysis, Investigation, Writing – original draft, Writing – review & editing. **Carla Novais:** Conceptualization, Formal analysis, Investigation, Writing – review & editing, Funding acquisition. **Mafalda Magalhães**: Investigation, Visualization. **Andreia Rebelo**: Investigation, Formal analysis, Visualization. **Sofia Ribeiro**: Investigation, Visualization. **Luísa Peixe:** Supervision, Funding acquisition, Writing – review & editing. **Ângela Novais:** Conceptualization, Methodology, Formal analysis, Writing –review & editing. **Patrícia Antunes:** Conceptualization, Methodology, Software, Formal analysis, Investigation, Supervision, Writing – original draft, Writing – review & editing, Funding acquisition, Project administration.

## REFERENCES

1. United Nations Environment Programme. 2023. Bracing for Superbugs: Strengthening environmental action in the One Health response to antimicrobial resistance. United Nations Environment Programme Geneva. https://www.unep.org/resources/superbugs/environmental-action.

2. World Health Organization. 2019. Critically Important Antimicrobials for Human Medicine 6th revision: Ranking of medically important antimicrobials for risk management of antimicrobial resistance due to non-human use. WHO Advisory Group on Integrated Surveillance of Antimicrobial Resistance (AGISAR). https://www.who.int/publications/i/item/9789241515528.

3. European Medicines Agency, Committee for Medicinal Products for Veterinary use, and Committee for Medicinal Products for Human Use. 2016. Updated advice on the use of colistin products in animals within the European Union: development of resistance and possible impact on human and animal health. https://www.ema.europa.eu/en/documents/scientific-guideline/updated-advice-use-colistin-products-animals-within-european-union-development-resistance-possible_en-0.pdf.

4. Jansen W, van Hout J, Wiegel J, Iatridou D, Chantziaras I, De Briyne N. 2022. Colistin Use in European Livestock: Veterinary Field Data on Trends and Perspectives for Further Reduction. Veterinary Sciences 9:650.

5. Olaitan AO, Dandachi I, Baron SA, Daoud Z, Morand S, Rolain J-M. 2021. Banning colistin in feed additives: a small step in the right direction. The Lancet Infectious Diseases 21:29–30.

6. European Medicines Agency, and Committee for Medicinal Products for Veterinary use. 2019. Advice on the designation of antimicrobials or groups of antimicrobials reserved for treatment of certain infections in humans - in relation to implementing measures under Article 37(5) of Regulation (EU) 2019/6 on veterinary medicinal products. https://www.ema.europa.eu/en/documents/regulatory-procedural-guideline/advice-designation-antimicrobials-groups-antimicrobials-reserved-treatment-certain-infections-humans/6-veterinary-medicinal-products_en.pdf.

7. European Medicines Agency, Committee for Medicinal Products for Veterinary use, and Committee for Medicinal Products for Human Use. 2019. Categorisation of antibiotics used in animals promotes responsible use to protect public and animal health: Answer to the request from the European Commission for updating the scientific advice on the impact on public health and animal health of the use of antibiotics in animals. https://www.ema.europa.eu/en/documents/press-release/categorisation-antibiotics-used-animals-promotes-responsible-use-protect-public-animal-health_en.pdf.

8. European Commission. 2020. Farm to Fork Strategy: For a fair, healthy and environmentally-friendly food system. https://food.ec.europa.eu/system/files/2020-05/f2f_action-plan_2020_strategy-info_en.pdf.

9. Regulation (EU) 2019/4. 2019. Regulation (EU) 2019/6 of the European Parliament and of the Council of 11 December 2018 on the manufacture, placing on the market and use of medicated feed, amending Regulation (EC) No 183/2005 of the European Parliament and of the Council and repealing Council Directive 90/167/EEC. Official Journal of the European Union. The European Parliament and the Council of the European Union. https://eur-lex.europa.eu/legal-content/EN/TXT/PDF/?uri=CELEX:32019R0004.

10. Regulation (EU) 2019/6. 2019. Regulation (EU) 2019/6 of the European Parliament and of the Council of 11 December 2018 on veterinary medicinal products and repealing Directive 2001/82/EC. Official Journal of the European Union. The European Parliament and the Council of the European Union. https://eur-lex.europa.eu/legal-content/EN/TXT/PDF/?uri=CELEX:32019R0006.

11. Rhouma M, Madec J-Y, Laxminarayan R. 2023. Colistin: from the shadows to a One Health approach for addressing antimicrobial resistance. International Journal of Antimicrobial Agents 61:106713.

12. Ribeiro S, Mourão J, Novais Â, Campos J, Peixe L, Antunes P. 2021. From farm to fork: Colistin voluntary withdrawal in Portuguese farms reflected in decreasing occurrence of *mcr-1-*carrying *Enterobacteriaceae* from chicken meat. Environmental Microbiology 23:7563–7577.

13. European Parliament, Augère-Granier M-L. 2019. The EU poultry meat and egg sector - Main features, challenges and prospects: in-depth analysis. Publications Office. https://data.europa.eu/doi/10.2861/33350.

14. Eurostat. 2021. Agricultural Production Livestock and Meat. https://ec.europa.eu/eurostat/statistics-explained/index.php?title=Agricultural_production_-_livestock_and_meat&oldid=427096#Meat_production.

15. Gadde U, Kim WH, Oh ST, Lillehoj HS. 2017. Alternatives to antibiotics for maximizing growth performance and feed efficiency in poultry: a review. Anim Health Res Rev 18:26– 45.

16. Betts JW, Hornsey M, La Ragione RM. 2018. Novel Antibacterials: Alternatives to Traditional Antibiotics, p. 123–169. In Advances in Microbial Physiology. Elsevier.

17. EMA Committee for Medicinal Products for Veterinary Use and EFSA Panel on Biological Hazards, Murphy D, Ricci A, Auce Z, Beechinor JG, Bergendahl H, Breathnach R, Bureš J, Duarte Da Silva JP, Hederová J, Hekman P, Ibrahim C, Kozhuharov E, Kulcsár G, Lander Persson E, Lenhardsson JM, Mačiulskis P, Malemis I, Markus-Cizelj L, Michaelidou-Patsia A, Nevalainen M, Pasquali P, Rouby J, Schefferlie J, Schlumbohm W, Schmit M, Spiteri S, Srčič S, Taban L, Tiirats T, Urbain B, Vestergaard E, Wachnik-Święcicka A, Weeks J, Zemann B, Allende A, Bolton D, Chemaly M, Fernandez Escamez PS, Girones R, Herman L, Koutsoumanis K, Lindqvist R, Nørrung B, Robertson L, Ru G, Sanaa M, Simmons M, Skandamis P, Snary E, Speybroeck N, Ter Kuile B, Wahlström H, Baptiste K, Catry B, Cocconcelli PS, Davies R, Ducrot C, Friis C, Jungersen G, More S, Muñoz Madero C, Sanders P, Bos M, Kunsagi Z, Torren Edo J, Brozzi R, Candiani D, Guerra B, Liebana E, Stella P, Threlfall J, Jukes H. 2017. EMA and EFSA Joint Scientific Opinion on measures to reduce the need to use antimicrobial agents in animal husbandry in the European Union, and the resulting impacts on food safety (RONAFA). EFSA J 15:e04666.

18. Broom LJ, Monteiro A, Piñon A. 2021. Recent Advances in Understanding the Influence of Zinc, Copper, and Manganese on the Gastrointestinal Environment of Pigs and Poultry. Animals 11:1276.

19. El Sabry MI, Stino FKR, El-Ghany WAA. 2021. Copper: benefits and risks for poultry, livestock, and fish production. Trop Anim Health Prod 53:487.

20. 20. EFSA Panel on Additives and Products or Substances used in Animal Feed. 2016. Revision of the currently authorised maximum copper content in complete feed. EFSA J 14:4563.

21. Jankowski J, Ognik K, Kozłowski K, Stępniowska A, Zduńczyk Z. 2019. Effect of Different Levels and Sources of Dietary Copper, Zinc and Manganese on the Performance and Immune and Redox Status of Turkeys. Animals 9:883.

22. Khatun A, Chowdhury S, Roy B, Dey B, Haque A, Chandran B. 2019. Comparative Effects of Inorganic and Three Forms of Organic Trace Minerals on Growth Performance, Carcass Traits, Immunity and Profitability of Broilers. J Adv Vet Anim Res 1:66–73.

23. 23. Tavares T, Mourão JL, Kay Z, Spring P, Vieira J, Gomes A, Vieira-Pinto M. 2013. The effect of replacing inorganic trace minerals with organic Bioplex® and Sel-Plex® on the performance and meat quality of broilers. J appl anim nutr 2:e10.

24. Rensing C, Moodley A, Cavaco LM, McDevitt SF. 2018. Resistance to Metals Used in Agricultural Production. Microbiol Spectr 6:6.2.20.

25. Anedda E, Farrell ML, Morris D, Burgess CM. 2023. Evaluating the impact of heavy metals on antimicrobial resistance in the primary food production environment: A scoping review. Environmental Pollution 320:121035.

26. Binsker U, Käsbohrer A, Hammerl JA. 2022. Global colistin use: a review of the emergence of resistant Enterobacterales and the impact on their genetic basis. FEMS Microbiology Reviews 46:fuab049.

27. Davis GS, Price LB. 2016. Recent Research Examining Links Among *Klebsiella pneumoniae* from Food, Food Animals, and Human Extraintestinal Infections. Curr Envir Health Rpt 3:128–135.

28. Wyres KL, Lam MMC, Holt KE. 2020. Population genomics of *Klebsiella pneumoniae*. Nat Rev Microbiol 18:344–359.

29. Wyres KL, Holt KE. 2016. *Klebsiella pneumoniae* Population Genomics and Antimicrobial-Resistant Clones. Trends in Microbiology 24:944–956.

30. Foster-Nyarko E, Cottingham H, Wick RR, Judd LM, Lam MMC, Wyres KL, Stanton TD, Tsang KK, David S, Aanensen DM, Brisse S, Holt KE. 2022. Nanopore-only assemblies for genomic surveillance of the global priority drug-resistant pathogen, Klebsiella pneumoniae. preprint. Genomics.

31. Martak D, Guther J, Verschuuren TD, Valot B, Conzelmann N, Bunk S, Riccio ME, Salamanca E, Meunier A, Henriot CP, Brossier CP, Bertrand X, Cooper BS, Harbarth S, Tacconelli E, Fluit AC, Rodriguez-Baño J, Kluytmans JAJW, Peter S, Hocquet D. 2022. Populations of extended-spectrum β-lactamase-producing *Escherichia coli* and *Klebsiella pneumoniae* are different in human-polluted environment and food items: a multicentre European study. Clinical Microbiology and Infection 28:447.e7-447.e14.

32. Schürch AC, Arredondo-Alonso S, Willems RJL, Goering RV. 2018. Whole genome sequencing options for bacterial strain typing and epidemiologic analysis based on single nucleotide polymorphism versus gene-by-gene–based approaches. Clinical Microbiology and Infection 24:350–354.

33. David S, Reuter S, Harris SR, Glasner C, Feltwell T, Argimon S, Abudahab K, Goater R, Giani T, Errico G, Aspbury M, Sjunnebo S, the EuSCAPE Working Group, Koraqi A, Lacej D, Apfalter P, Hartl R, Glupczynski Y, Huang T-D, Strateva T, Marteva-Proevska Y, Andrasevic AT, Butic I, Pieridou-Bagatzouni D, Maikanti-Charalampous P, Hrabak J, Zemlickova H, Hammerum A, Jakobsen L, Ivanova M, Pavelkovich A, Jalava J, Österblad M, Dortet L, Vaux S, Kaase M, Gatermann SG, Vatopoulos A, Tryfinopoulou K, Tóth Á, Jánvári L, Boo TW, McGrath E, Carmeli Y, Adler A, Pantosti A, Monaco M, Raka L, Kurti A, Balode A, Saule M, Miciuleviciene J, Mierauskaite A, Perrin-Weniger M, Reichert P, Nestorova N, Debattista S, Mijovic G, Lopicic M, Samuelsen Ø, Haldorsen B, Zabicka D, Literacka E, Caniça M, Manageiro V, Kaftandzieva A, Trajkovska-Dokic E, Damian M, Lixandru B, Jelesic Z, Trudic A, Niks M, Schreterova E, Pirs M, Cerar T, Oteo J, Aracil B, Giske C, Sjöström K, Gür D, Cakar A, Woodford N, Hopkins K, Wiuff C, Brown DJ, the ESGEM Study Group, Feil EJ, Rossolini GM, Aanensen DM, Grundmann H. 2019. Epidemic of carbapenem-resistant Klebsiella pneumoniae in Europe is driven by nosocomial spread. Nat Microbiol 4:1919–1929.

34. Daehre K, Projahn M, Friese A, Semmler T, Guenther S, Roesler UH. 2018. ESBL-Producing *Klebsiella pneumoniae* in the Broiler Production Chain and the First Description of ST3128. Front Microbiol 9:2302.

35. Zhai R, Fu B, Shi X, Sun C, Liu Z, Wang S, Shen Z, Walsh TR, Cai C, Wang Y, Wu C. 2020. Contaminated in-house environment contributes to the persistence and transmission of NDM-producing bacteria in a Chinese poultry farm. Environment International 139:105715.

36. Kers JG, Velkers FC, Fischer EAJ, Hermes GDA, Stegeman JA, Smidt H. 2018. Host and Environmental Factors Affecting the Intestinal Microbiota in Chickens. Front Microbiol 9:235.

37. Marmion M, Ferone MT, Whyte P, Scannell AGM. 2021. The changing microbiome of poultry meat; from farm to fridge. Food Microbiology 99:103823.

38. Franklin-Alming FV, Kaspersen H, Hetland MAK, Bakksjø R-J, Nesse LL, Leangapichart T, Löhr IH, Telke AA, Sunde M. 2021. Exploring *Klebsiella pneumoniae* in Healthy Poultry Reveals High Genetic Diversity, Good Biofilm-Forming Abilities and Higher Prevalence in Turkeys Than Broilers. Front Microbiol 12:725414.

39. Davis GS, Waits K, Nordstrom L, Weaver B, Aziz M, Gauld L, Grande H, Bigler R, Horwinski J, Porter S, Stegger M, Johnson JR, Liu CM, Price LB. 2015. Intermingled *Klebsiella pneumoniae* Populations Between Retail Meats and Human Urinary Tract Infections. Clin Infect Dis 61:892–899.

40. Rodrigues C, Hauser K, Cahill N, Ligowska-Marzęta M, Centorotola G, Cornacchia A, Garcia Fierro R, Haenni M, Nielsen EM, Piveteau P, Barbier E, Morris D, Pomilio F, Brisse S. 2022. High Prevalence of *Klebsiella pneumoniae* in European Food Products: a Multicentric Study Comparing Culture and Molecular Detection Methods. Microbiol Spectr 10:e02376–21.

41. Projahn M, von Tippelskirch P, Semmler T, Guenther S, Alter T, Roesler U. 2019. Contamination of chicken meat with extended-spectrum beta-lactamase producing-*Klebsiella pneumoniae* and *Escherichia coli* during scalding and defeathering of broiler carcasses. Food Microbiology 77:185–191.

42. Cannatelli A, Giani T, D’Andrea MM, Di Pilato V, Arena F, Conte V, Tryfinopoulou K, Vatopoulos A, Rossolini GM. 2014. MgrB Inactivation Is a Common Mechanism of Colistin Resistance in KPC-Producing *Klebsiella pneumoniae* of Clinical Origin. Antimicrob Agents Chemother 58:5696–5703.

43. Elias R, Spadar A, Phelan J, Melo-Cristino J, Lito L, Pinto M, Gonçalves L, Campino S, Clark TG, Duarte A, Perdigão J. 2022. A phylogenomic approach for the analysis of colistin resistance-associated genes in *Klebsiella pneumoniae*, its mutational diversity and implications for phenotypic resistance. International Journal of Antimicrobial Agents 59:106581.

44. Janssen AB, Doorduijn DJ, Mills G, Rogers MRC, Bonten MJM, Rooijakkers SHM, Willems RJL, Bengoechea JA, van Schaik W. 2020. Evolution of Colistin Resistance in the *Klebsiella pneumoniae* Complex Follows Multiple Evolutionary Trajectories with Variable Effects on Fitness and Virulence Characteristics. Antimicrob Agents Chemother 65:e01958–20.

45. Jayol A, Nordmann P, Lehours P, Poirel L, Dubois V. 2018. Comparison of methods for detection of plasmid-mediated and chromosomally encoded colistin resistance in Enterobacteriaceae. Clin Microbiol Infect 24:175–179.

46. Malli E, Florou Z, Tsilipounidaki K, Voulgaridi I, Stefos A, Xitsas S, Papagiannitsis CC, Petinaki E. 2018. Evaluation of rapid polymyxin NP test to detect colistin-resistant *Klebsiella pneumoniae* isolated in a tertiary Greek hospital. J Microbiol Methods 153:35– 39.

47. Mathur P, Veeraraghavan B, Devanga Ragupathi NK, Inbanathan FY, Khurana S, Bhardwaj N, Kumar S, Sagar S, Gupta A. 2018. Multiple mutations in lipid-A modification pathway & novel *fosA* variants in colistin-resistant *Klebsiella pneumoniae*. Future Sci OA 4:FSO319.

48. Ngbede EO, Adekanmbi F, Poudel A, Kalalah A, Kelly P, Yang Y, Adamu AM, Daniel ST, Adikwu AA, Akwuobu CA, Abba PO, Mamfe LM, Maurice NA, Adah MI, Lockyear O, Butaye P, Wang C. 2021. Concurrent Resistance to Carbapenem and Colistin Among Enterobacteriaceae Recovered From Human and Animal Sources in Nigeria Is Associated With Multiple Genetic Mechanisms. Front Microbiol 12:740348.

49. Nordmann P, Jayol A, Poirel L. 2016. Rapid Detection of Polymyxin Resistance in Enterobacteriaceae. Emerg Infect Dis 22:1038–1043.

50. Tietgen M, Sedlaczek L, Higgins PG, Kaspar H, Ewers C, Göttig S. 2022. Colistin Resistance Mechanisms in Human and Veterinary *Klebsiella pneumoniae* Isolates. Antibiotics (Basel) 11:1672.

51. Furlan JPR, Stehling EG, Savazzi EA, Peixe L, Novais Â. 2022. High occurrence of colistin- and multidrug-resistant strains carrying *mcr*-1 or an underestimated *mcr*-1.26 allelic variant along a large Brazilian river. Journal of Global Antimicrobial Resistance 30:127–129.

52. Bray AS, Smith RD, Hudson AW, Hernandez GE, Young TM, George HE, Ernst RK, Zafar MA. 2022. MgrB-Dependent Colistin Resistance in *Klebsiella pneumoniae* Is Associated with an Increase in Host-to-Host Transmission. mBio 13:e03595–21.

53. Wand ME, Bock LJ, Bonney LC, Sutton JM. 2017. Mechanisms of Increased Resistance to Chlorhexidine and Cross-Resistance to Colistin following Exposure of *Klebsiella pneumoniae* Clinical Isolates to Chlorhexidine. Antimicrob Agents Chemother 61:e01162–16.

54. Zhang Y, Zhao Y, Xu C, Zhang X, Li J, Dong G, Cao J, Zhou T. 2019. Chlorhexidine exposure of clinical *Klebsiella pneumoniae* strains leads to acquired resistance to this disinfectant and to colistin. International Journal of Antimicrobial Agents 53:864–867.

55. Cannatelli A, Santos-Lopez A, Giani T, Gonzalez-Zorn B, Rossolini GM. 2015. Polymyxin Resistance Caused by *mgrB* Inactivation Is Not Associated with Significant Biological Cost in *Klebsiella pneumoniae*. Antimicrob Agents Chemother 59:2898–2900.

56. Zhu X-Q, Liu Y-Y, Wu R, Xun H, Sun J, Li J, Feng Y, Liu J-H. 2021. Impact of *mcr*-1 on the Development of High Level Colistin Resistance in *Klebsiella pneumoniae* and *Escherichia coli*. Front Microbiol 12:666782.

57. Tu Z, Gu J, Zhang H, Liu J, Shui J, Zhang A. 2021. Withdrawal of Colistin Reduces Incidence of *mcr*-1-Harboring IncX4-Type Plasmids but Has Limited Effects on Unrelated Antibiotic Resistance. Pathogens 10:1019.

58. 58. Zhang Q, Lv L, Huang X, Huang Y, Zhuang Z, Lu J, Liu E, Wan M, Xun H, Zhang Z, Huang J, Song Q, Zhuo C, Liu J-H. 2019. Rapid Increase in Carbapenemase-Producing Enterobacteriaceae in Retail Meat Driven by the Spread of the *bla _NDM-5_*-Carrying IncX3 Plasmid in China from 2016 to 2018. Antimicrob Agents Chemother 63:e00573–19.

59. Savin M, Bierbaum G, Schmithausen RM, Heinemann C, Kreyenschmidt J, Schmoger S, Akbaba I, Käsbohrer A, Hammerl JA. 2022. Slaughterhouse wastewater as a reservoir for extended-spectrum β-lactamase (ESBL)-producing, and colistin-resistant *Klebsiella* spp. and their impact in a “One Health” perspective. Science of The Total Environment 804:150000.

60. Thorpe HA, Booton R, Kallonen T, Gibbon MJ, Couto N, Passet V, López-Fernández S, Rodrigues C, Matthews L, Mitchell S, Reeve R, David S, Merla C, Corbella M, Ferrari C, Comandatore F, Marone P, Brisse S, Sassera D, Corander J, Feil EJ. 2022. A large-scale genomic snapshot of *Klebsiella* spp. isolates in Northern Italy reveals limited transmission between clinical and non-clinical settings. Nat Microbiol 7:2054–2067.

61. Liu C, Li G, Qin X, Xu Y, Wang J, Wu G, Feng H, Ye J, Zhu C, Li X, Zheng X. 2022. Profiles of antibiotic- and heavy metal-related resistance genes in animal manure revealed using a metagenomic analysis. Ecotoxicology and Environmental Safety 239:113655.

62. Mazhar SH, Li X, Rashid A, Su J, Xu J, Brejnrod AD, Su J-Q, Wu Y, Zhu Y-G, Zhou SG, Feng R, Rensing C. 2021. Co-selection of antibiotic resistance genes, and mobile genetic elements in the presence of heavy metals in poultry farm environments. Science of The Total Environment 755:142702.

63. Mourão J, Marçal S, Ramos P, Campos J, Machado J, Peixe L, Novais C, Antunes P. 2016. Tolerance to multiple metal stressors in emerging non-typhoidal MDR *Salmonella* serotypes: A relevant role for copper in anaerobic conditions. Journal of Antimicrobial Chemotherapy 71:2147–2157.

64. Yan X, Yang J, Wang Q, Lin S. 2021. Transcriptomic analysis reveals resistance mechanisms of *Klebsiella michiganensis* to copper toxicity under acidic conditions. Ecotoxicology and Environmental Safety 211:111919.

65. Gullberg E, Albrecht LM, Karlsson C, Sandegren L, Andersson DI. 2014. Selection of a Multidrug Resistance Plasmid by Sublethal Levels of Antibiotics and Heavy Metals. mBio 5:e01918–14.

66. Li X, Gu AZ, Zhang Y, Xie B, Li D, Chen J. 2019. Sub-lethal concentrations of heavy metals induce antibiotic resistance via mutagenesis. Journal of Hazardous Materials 369:9–16.

67. Håkonsholm F, Hetland MAK, Svanevik CS, Sundsfjord A, Lunestad BT, Marathe NP. 2020. Antibiotic Sensitivity Screening of *Klebsiella* spp. and *Raoultella* spp. Isolated from Marine Bivalve Molluscs Reveal Presence of CTX-M-Producing K. pneumoniae. Microorganisms 8:1909.

68. Rodrigues C, Machado E, Ramos H, Peixe L, Novais Â. 2014. Expansion of ESBL-producing Klebsiella pneumoniae in hospitalized patients: A successful story of international clones (ST15, ST147, ST336) and epidemic plasmids (IncR, IncFIIK). International Journal of Medical Microbiology 304:1100–1108.

69. Bai S, Yu Y, Kuang X, Li X, Wang M, Sun R, Sun J, Liu Y, Liao X. 2022. Molecular Characteristics of Antimicrobial Resistance and Virulence in *Klebsiella pneumoniae* Strains Isolated from Goose Farms in Hainan, China. Appl Environ Microbiol 88:e02457–21.

70. Matlock W, Chau KK, AbuOun M, Stubberfield E, Barker L, Kavanagh J, Pickford H, Gilson D, Smith RP, Gweon HS, Hoosdally SJ, Swann J, Sebra R, Bailey MJ, Peto TEA, Crook DW, Anjum MF, Read DS, Walker AS, Stoesser N, Shaw LP, REHAB consortium, AbuOun M, Anjum MF, Bailey MJ, Brett H, Bowes MJ, Chau KK, Crook DW, de Maio N, Duggett N, Wilson DJ, Gilson D, Gweon HS, Hubbard A, Hoosdally SJ, Matlock W, Kavanagh J, Jones H, Peto TEA, Read DS, Sebra R, Shaw LP, Sheppard AE, Smith RP, Stubberfield E, Stoesser N, Swann J, Walker AS, Woodford N. 2021. Genomic network analysis of environmental and livestock F-type plasmid populations. ISME J 15:2322– 2335.

71. Rocha J, Ferreira C, Mil-Homens D, Busquets A, Fialho AM, Henriques I, Gomila M, Manaia CM. 2022. Third generation cephalosporin-resistant *Klebsiella pneumoniae* thriving in patients and in wastewater: what do they have in common? BMC Genomics 23:72.

72. Rodrigues C, Lanza VF, Peixe L, Coque TM, Novais Â. 2022. Phylogenomics of globally spread Clonal Groups 14 and 15 of *Klebsiella pneumoniae*. preprint. Genomics.

73. Spadar A, Perdigão J, Campino S, Clark TG. 2023. Large-scale genomic analysis of global *Klebsiella pneumoniae* plasmids reveals multiple simultaneous clusters of carbapenem-resistant hypervirulent strains. Genome Med 15:3.

74. Ludden C, Moradigaravand D, Jamrozy D, Gouliouris T, Blane B, Naydenova P, Hernandez-Garcia J, Wood P, Hadjirin N, Radakovic M, Crawley C, Brown NM, Holmes M, Parkhill J, Peacock SJ. 2020. A One Health Study of the Genetic Relatedness of *Klebsiella pneumoniae* and Their Mobile Elements in the East of England. Clinical Infectious Diseases 70:219–226.

75. Savin M, Bierbaum G, Blau K, Parcina M, Sib E, Smalla K, Schmithausen R, Heinemann C, Hammerl JA, Kreyenschmidt J. 2020. Colistin-Resistant Enterobacteriaceae Isolated From Process Waters and Wastewater From German Poultry and Pig Slaughterhouses. Front Microbiol 11:575391.

76. Rodrigues C, Sousa C, Lopes JA, Novais Â, Peixe L. 2020. A Front Line on *Klebsiella pneumoniae* Capsular Polysaccharide Knowledge: Fourier Transform Infrared Spectroscopy as an Accurate and Fast Typing Tool. mSystems 5:e00386–19.

77. Chen L, Mathema B, Pitout JDD, DeLeo FR, Kreiswirth BN. 2014. Epidemic *Klebsiella pneumoniae* ST258 Is a Hybrid Strain. mBio 5:e01355–14.

78. Mostowy RJ, Holt KE. 2018. Diversity-Generating Machines: Genetics of Bacterial Sugar-Coating. Trends in Microbiology 26:1008–1021.

79. Patro LPP, Rathinavelan T. 2019. Targeting the Sugary Armor of *Klebsiella* Species. Front Cell Infect Microbiol 9:367.

80. 80. Association of Poultry Processors and Poultry Trade in the EU Countries Ltd. 2016. Comparison of the Regulatory Framework and Key Practices in the Poultry Meat Supply Chain in the EU and USA. https://britishpoultry.org.uk/identity-cms/wp-content/uploads/2018/05/2016-ADAS-EU-US-comparison.pdf.

81. Bialek-Davenet S, Criscuolo A, Ailloud F, Passet V, Nicolas-Chanoine M-H, Decré D, Brisse S. 2014. Development of a multiplex PCR assay for identification of *Klebsiella pneumoniae* hypervirulent clones of capsular serotype K2. Journal of Medical Microbiology 63:1608–1614.

82. Borowiak M, Baumann B, Fischer J, Thomas K, Deneke C, Hammerl JA, Szabo I, Malorny B. 2020. Development of a Novel *mcr*-6 to *mcr*-9 Multiplex PCR and Assessment of *mcr*-1 to *mcr*-9 Occurrence in Colistin-Resistant *Salmonella enterica* Isolates From Environment, Feed, Animals and Food (2011–2018) in Germany. Front Microbiol 11:80.

83. 83. Rebelo AR, Bortolaia V, Kjeldgaard JS, Pedersen SK, Leekitcharoenphon P, Hansen IM, Guerra B, Malorny B, Borowiak M, Hammerl JA, Battisti A, Franco A, Alba P, Perrin-Guyomard A, Granier SA, De Frutos Escobar C, Malhotra-Kumar S, Villa L, Carattoli A, Hendriksen RS. 2018. Multiplex PCR for detection of plasmid-mediated colistin resistance determinants, *mcr*-1, *mcr*-2, *mcr*-3, *mcr*-4 and *mcr*-5 for surveillance purposes. Eurosurveillance 23:17-00672.

84. 84. European Committee of Antimicrobial Susceptibility Testing. 2016. Recommendations for MIC determination of colistin (polymyxin E) - As recommended by the joint CLSI-EUCAST Polymyxin Breakpoints Working Group.

85. Sousa C, Novais Â, Magalhães A, Lopes J, Peixe L. 2013. Diverse high-risk B2 and D *Escherichia coli* clones depicted by Fourier Transform Infrared Spectroscopy. Sci Rep 3:3278.

86. Brisse S, Passet V, Haugaard AB, Babosan A, Kassis-Chikhani N, Struve C, Decré D. 2013. wzi Gene Sequencing, a Rapid Method for Determination of Capsular Type for *Klebsiella* Strains. J Clin Microbiol 51:4073–4078.

87. 87. European Committee on Antimicrobial Susceptibility Testing. 2022. Breakpoint tables for interpretation of MICs and zone diameters. Version 12.0.

88. 88. Clinical and Laboratory Standards Institute. 2022. Performance Standards for Antimicrobial Susceptibility Testing, 32nd Edition; CLSI Document M100.

89. Andrews S. 2010. FastQC: A quality control tool for high throughput sequence data (online).

90. Ewels P, Magnusson M, Lundin S, Käller M. 2016. MultiQC: summarize analysis results for multiple tools and samples in a single report. Bioinformatics 32:3047–3048.

91. Bankevich A, Nurk S, Antipov D, Gurevich AA, Dvorkin M, Kulikov AS, Lesin VM, Nikolenko SI, Pham S, Prjibelski AD, Pyshkin AV, Sirotkin AV, Vyahhi N, Tesler G, Alekseyev MA, Pevzner PA. 2012. SPAdes: A New Genome Assembly Algorithm and Its Applications to Single-Cell Sequencing. Journal of Computational Biology 19:455–477.

92. Gurevich A, Saveliev V, Vyahhi N, Tesler G. 2013. QUAST: quality assessment tool for genome assemblies. Bioinformatics 29:1072–1075.

93. Parks DH, Imelfort M, Skennerton CT, Hugenholtz P, Tyson GW. 2015. CheckM: assessing the quality of microbial genomes recovered from isolates, single cells, and metagenomes. Genome Res 25:1043–1055.

94. Aziz RK, Bartels D, Best AA, DeJongh M, Disz T, Edwards RA, Formsma K, Gerdes S, Glass EM, Kubal M, Meyer F, Olsen GJ, Olson R, Osterman AL, Overbeek RA, McNeil LK, Paarmann D, Paczian T, Parrello B, Pusch GD, Reich C, Stevens R, Vassieva O, Vonstein V, Wilke A, Zagnitko O. 2008. The RAST Server: Rapid Annotations using Subsystems Technology. BMC Genomics 9:75.

95. Diancourt L, Passet V, Verhoef J, Grimont PAD, Brisse S. 2005. Multilocus Sequence Typing of *Klebsiella pneumoniae* Nosocomial Isolates. J Clin Microbiol 43:4178–4182.

96. Argimón S, David S, Underwood A, Abrudan M, Wheeler NE, Kekre M, Abudahab K, Yeats CA, Goater R, Taylor B, Harste H, Muddyman D, Feil EJ, Brisse S, Holt K, Donado-Godoy P, Ravikumar KL, Okeke IN, Carlos C, Aanensen DM, NIHR Global Health Research Unit on Genomic Surveillance of Antimicrobial Resistance, Fabian Bernal J, Arevalo A, Fernanda Valencia M, Osma Castro ECD, Nagaraj G, Shamanna V, Govindan V, Prabhu A, Sravani D, Shincy MR, Rose S, Ravishankar KN, Oaikhena AO, Afolayan AO, Ajiboye JJ, Ewomazino Odih E, Lagrada ML, Macaranas PKV, Olorosa AM, Gayeta JM, Masim MAL, Herrera EM, Molloy A, Stelling J. 2021. Rapid Genomic Characterization and Global Surveillance of *Klebsiella* Using Pathogenwatch. Clinical Infectious Diseases 73:S325–S335.

97. Letunic I, Bork P. 2021. Interactive Tree Of Life (iTOL) v5: an online tool for phylogenetic tree display and annotation. Nucleic Acids Research 49:W293–W296.

98. Formosa C, Herold M, Vidaillac C, Duval RE, Dague E. 2015. Unravelling of a mechanism of resistance to colistin in *Klebsiella pneumoniae* using atomic force microscopy. Journal of Antimicrobial Chemotherapy 70:2261–2270.

99. Choi Y, Chan AP. 2015. PROVEAN web server: a tool to predict the functional effect of amino acid substitutions and indels. Bioinformatics 31:2745–2747.

100. Lam MMC, Wick RR, Watts SC, Cerdeira LT, Wyres KL, Holt KE. 2021. A genomic surveillance framework and genotyping tool for *Klebsiella pneumoniae* and its related species complex. Nat Commun 12:4188.

101. Gato E, Vázquez-Ucha JC, Rumbo-Feal S, Álvarez-Fraga L, Vallejo JA, Martínez-Guitián M, Beceiro A, Ramos Vivas J, Sola Campoy PJ, Pérez-Vázquez M, Oteo Iglesias J, Rodiño-Janeiro BK, Romero A, Poza M, Bou G, Pérez A. 2020. Kpi, a chaperone-usher pili system associated with the worldwide-disseminated high-risk clone *Klebsiella pneumoniae* ST-15. Proc Natl Acad Sci USA 117:17249–17259.

102. Feldgarden M, Brover V, Gonzalez-Escalona N, Frye JG, Haendiges J, Haft DH, Hoffmann M, Pettengill JB, Prasad AB, Tillman GE, Tyson GH, Klimke W. 2021. AMRFinderPlus and the Reference Gene Catalog facilitate examination of the genomic links among antimicrobial resistance, stress response, and virulence. Sci Rep 11:12728.

103. Camacho C, Coulouris G, Avagyan V, Ma N, Papadopoulos J, Bealer K, Madden TL. 2009. BLAST+: architecture and applications. BMC Bioinformatics 10:421.

104. 104. Carattoli A, Zankari E, García-Fernández A, Voldby Larsen M, Lund O, Villa L, Møller Aarestrup F, Hasman H. 2014. *In Silico* Detection and Typing of Plasmids using PlasmidFinder and Plasmid Multilocus Sequence Typing. Antimicrob Agents Chemother 58:3895–3903.

105. Villa L, García-Fernández A, Fortini D, Carattoli A. 2010. Replicon sequence typing of IncF plasmids carrying virulence and resistance determinants. Journal of Antimicrobial Chemotherapy 65:2518–2529.

106. Robertson J, Bessonov K, Schonfeld J, Nash JHE. 2020. Universal whole-sequence-based plasmid typing and its utility to prediction of host range and epidemiological surveillance. Microbial Genomics 6:mgen000435.

107. Robertson J, Nash JHE. 2018. MOB-suite: software tools for clustering, reconstruction and typing of plasmids from draft assemblies. Microbial Genomics 4:e000206.

