## Supplementary material for "From farm to fork: persistence of clinically-relevant multidrug-resistant and copper-tolerant *Klebsiella pneumoniae* long after colistin withdrawal in poultry production": Fig. S1, Fig. S2, Fig. S3 and Fig. S4


**Running Head:** Colistin-resistant *K. pneumoniae* in chicken production

*Joana Mourão, Carla Novais, Luísa Peixe and Patrícia Antunes are active members of the European Society of Clinical Microbiology and Infectious Diseases (ESCMID) Food and Water-borne Infection Study Group (EFWISG).

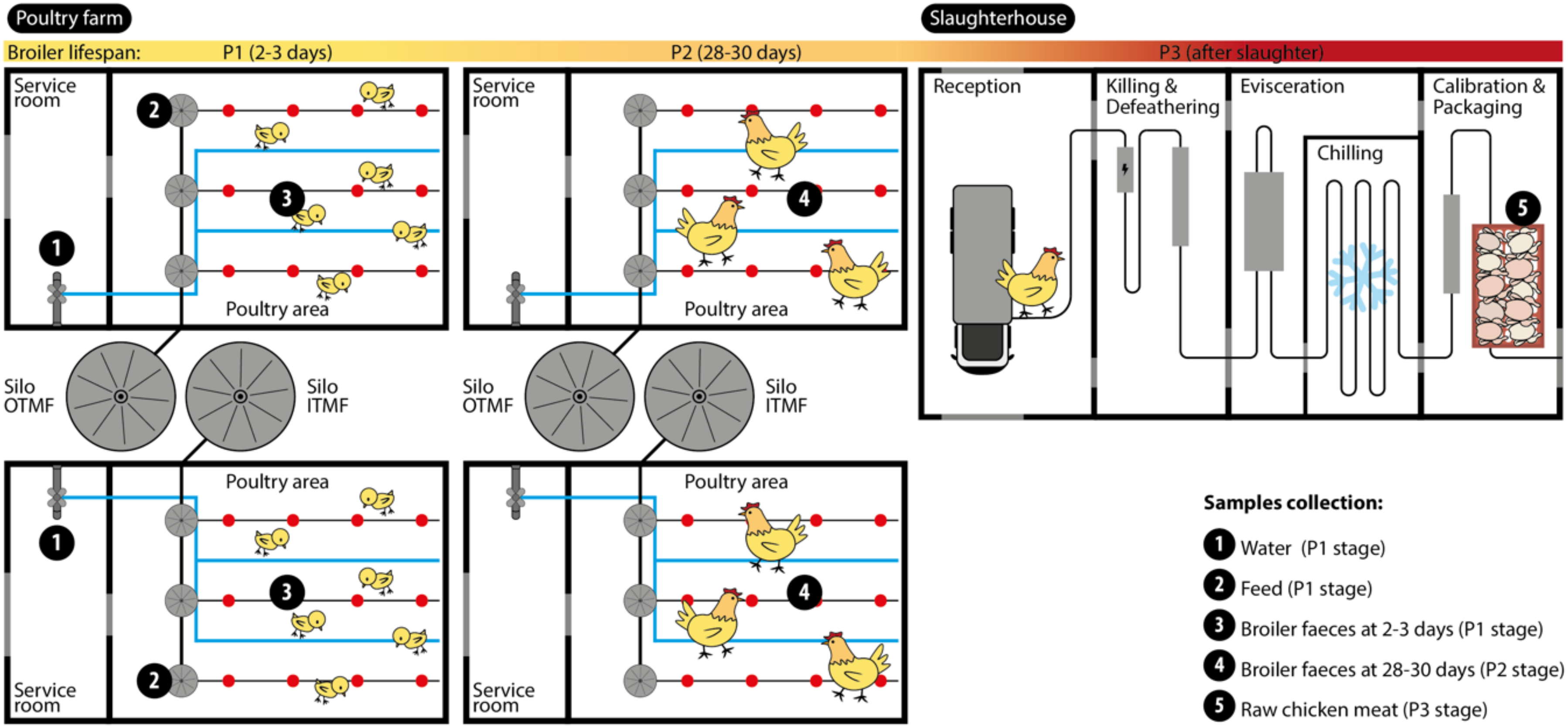


**Fig. S1.** Sampling strategy at the chicken farm and slaughterhouse processing plant. Sample collection points are indicated by numbers.

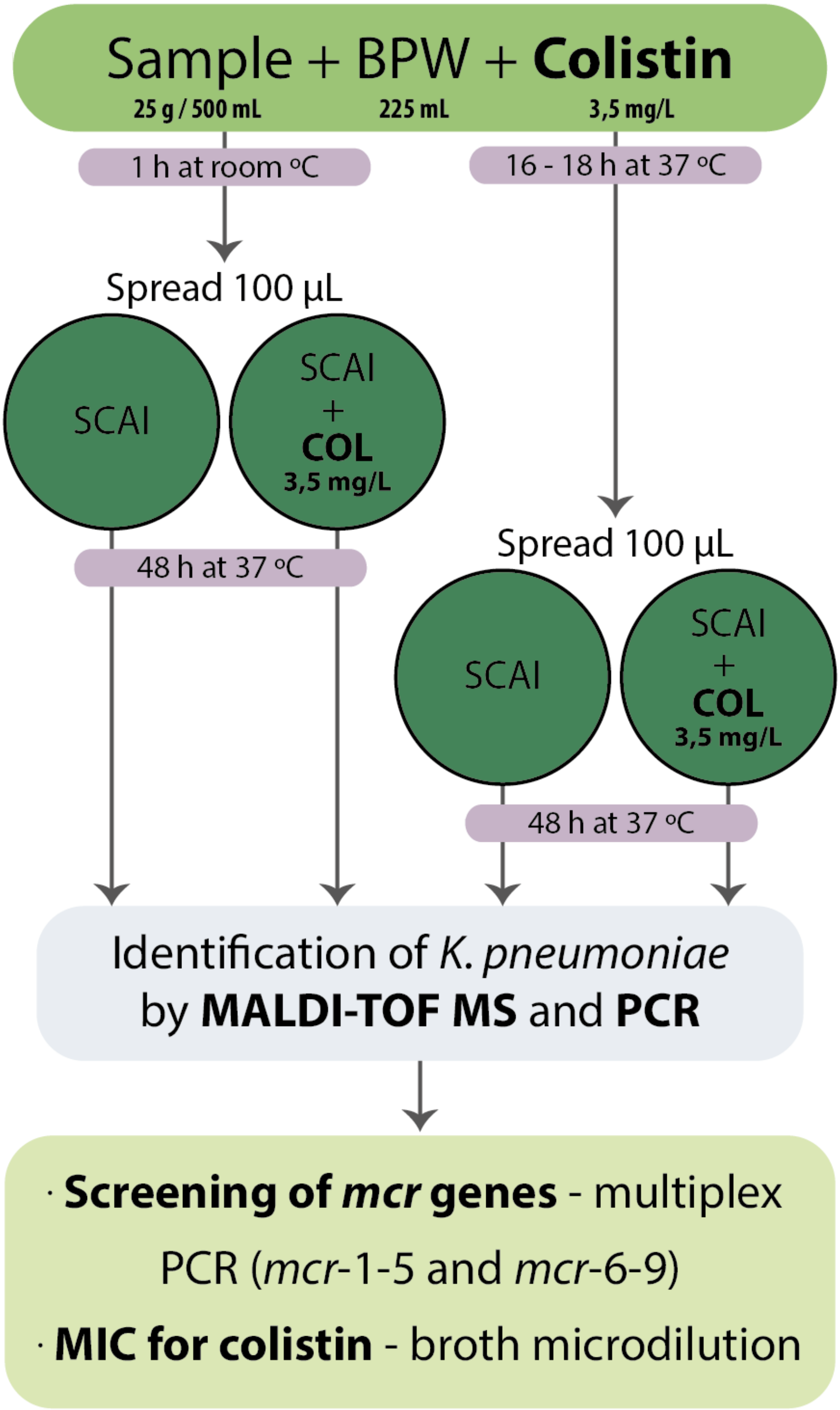


**Fig. S2.** Workflow for screening and counts of *K. pneumoniae and* colistin-resistant *K. pneumoniae*. BPW, buffered peptone water; COL, colistin.

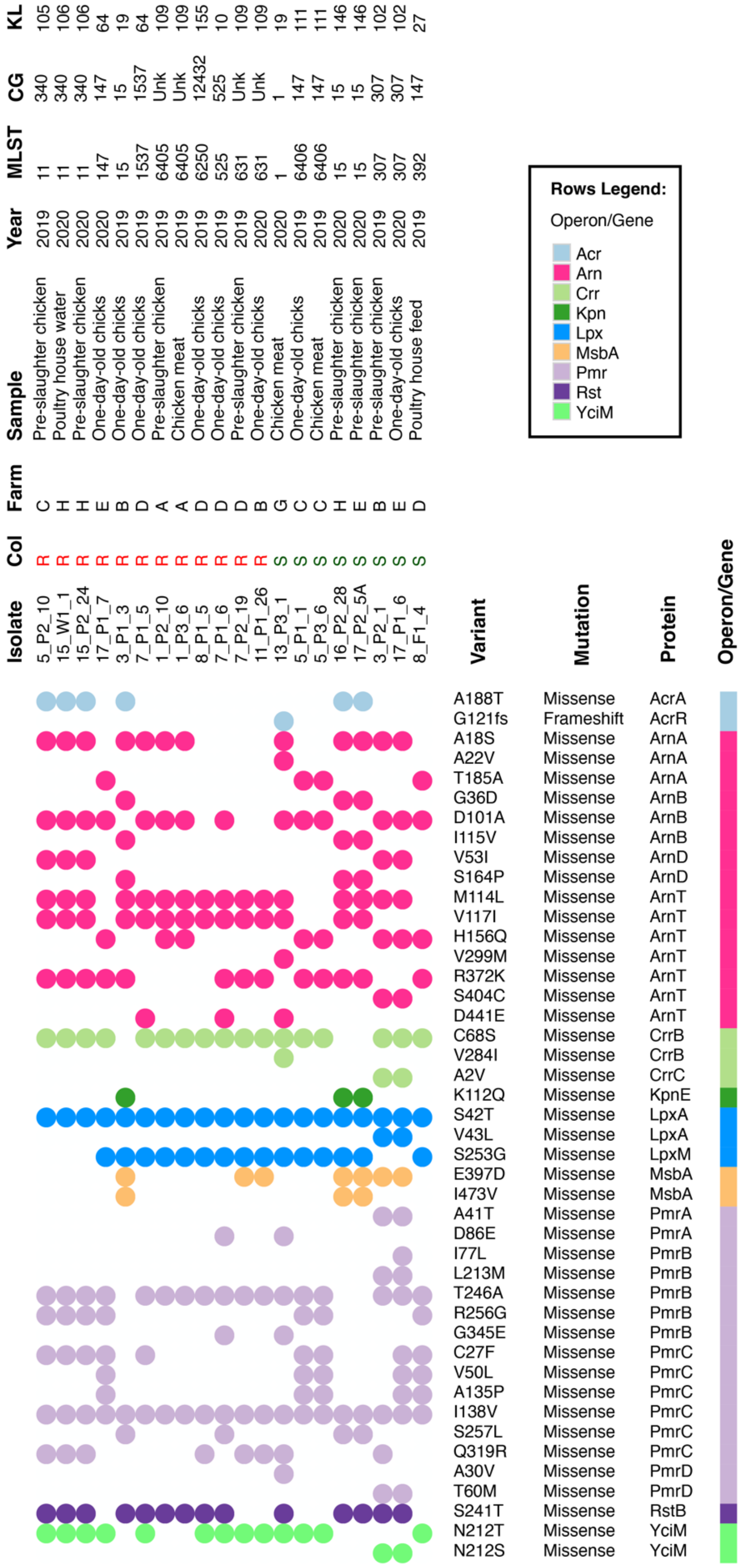
Isolate
Col
Farm
Sample
Year
MLST
CG
KL
5_P2_10 R C Pre-slaughter chicken 2019 11 340 105
15_W1_1 R H Poultry house water 2020 11 340 106
15_P2_24 R H Pre-slaughter chicken 2020 11 340 106
17_P1_7 R E One-day-old chicks 2020 147 147 64
3_P1_3 R B One-day-old chicks 2019 15 15 19
7_P1_5 R D One-day-old chicks 2019 1537 1537 64
1_P2_10 R A Pre-slaughter chicken 2019 6405 Unk 109
1_P3_6 R A Chicken meat 2019 6405 Unk 109
8_P1_5 R D One-day-old chicks 2019 6250 12432 155
7_P1_6 R D One-day-old chicks 2019 525 525 10
7_P2_19 R D Pre-slaughter chicken 2019 631 Unk 109
11_P1_26 R B One-day-old chicks 2020 631 Unk 109
13_P3_1 S G Chicken meat 2020 1 1 19
5_P1_1 S C One-day-old chicks 2019 6406 147 111
5_P3_6 S C Chicken meat 2019 6406 147 111
16_P2_28 S H Pre-slaughter chicken 2020 15 15 146
17_P2_5A S E Pre-slaughter chicken 2020 15 15 146
3_P2_1 S B Pre-slaughter chicken 2019 307 307 102
17_P1_6 S E One-day-old chicks 2020 307 307 102
8_F1_4 S D Poultry house feed 2019 392 147 27
Rows Legend:
Operon/Gene
Acr
Arn
Crr
Kpn
Lpx
MsbA
Pmr
Rst
YciM
Variant Mutation Protein Operon/Gene
A188T Missense AcrA
G121fs Frameshift AcrR
A18S Missense ArnA
A22V Missense ArnA
T185A Missense ArnA
G36D Missense ArnB
D101A Missense ArnB
I115V Missense ArnB
V53I Missense ArnD
S164P Missense ArnD
M114L Missense ArnT
V117I Missense ArnT
H156Q Missense ArnT
V299M Missense ArnT
R372K Missense ArnT
S404C Missense ArnT
D441E Missense ArnT
C68S Missense CrrB
V284I Missense CrrB
A2V Missense CrrC
K112Q Missense KpnE
S42T Missense LpxA
V43L Missense LpxA
S253G Missense LpxM
E397D Missense MsbA
I473V Missense MsbA
A41T Missense PmrA
D86E Missense PmrA
I77L Missense PmrB
L213M Missense PmrB
T246A Missense PmrB
R256G Missense PmrB
G345E Missense PmrB
C27F Missense PmrC
V50L Missense PmrC
A135P Missense PmrC
I138V Missense PmrC
S257L Missense PmrC
Q319R Missense PmrC
A30V Missense PmrD
T60M Missense PmrD
S241T Missense RstB
N212T Missense YciM
N212S Missense YciM

**Fig. S3.** Heatmap generated using Morpheus online web-server (https://software.broadinstitute.org/morpheus/) representing the distribution of colistin-associated mutations among the sequenced *Klebsiella pneumoniae* genomes. Coloured squares indicate the presence of a specific mutation, while each colour represents the genes that are part of the same operon. *Klebsiella pneumoniae* isolates were grouped into colistin-susceptible when MIC = <2 $\mu$g/mL or colistin-resistant when MIC = ≥2 $\mu$g/mL. When required, figures were minimally edited manually using Adobe Illustrator v25.3.1. Detailed resistance mechanisms: *acrA* – efflux pump; *acrR* – a transcriptional regulator of *acrAB* operon; *arnABDT* - lipid A modification with L-Ara4N addition; *crrB* – lipid A modification by upregulation of *pmrAB*/activation of the glycosyltransferase; *crrC* – connector protein; *kpnE* – efflux pump; *lpxA* – inactivation of lipid A biosynthesis abolishing LPS synthesis; *lpxM* – increased lipid A acylation; *msbA* - ABC transporter of lipid A; *pmrAB* - activation of LPS-modifying operation in the two-component systems; *pmrC* - lipid A modification with PEtn; *pmrD* – connector protein; *mgrB* - inactivation of negative feedback regulator of the PhoP/PhoQ system; *rstB* - sensor protein; *yciM* - regulation of LPS biosynthesis. Abbreviations: A, alanine; C, cysteine; D, aspartate; E, glutamate; F, phenylalanine; G, glycine; H, histidine; I, isoleucine; K, lysine; L, leucine; M, methionine (start codon); N, asparagine; P, proline; Q, glutamine; R, arginine; S, serine; T, threonine; V, valine; Unk, unknown.

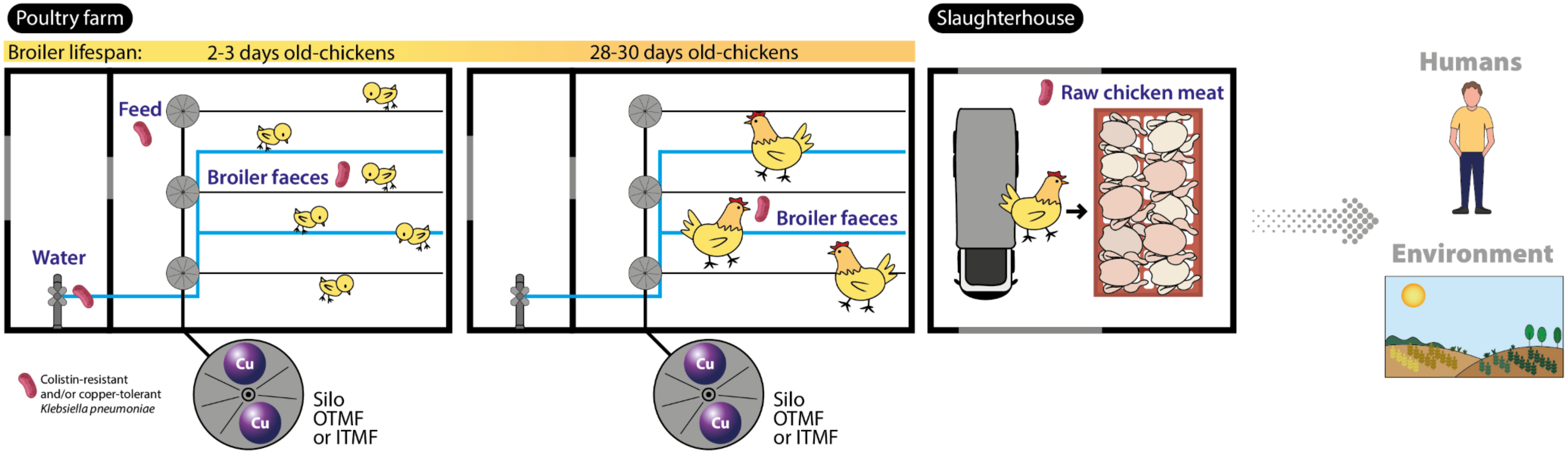


**Fig. S4.** Graphical abstract
